# Transcriptomics-informed pharmacology identifies epigenetic and cell cycle regulators as enhancers of AAV production

**DOI:** 10.1101/2024.06.14.599118

**Authors:** Joshua Tworig, Francis Grafton, Markus Hörer, Christopher A. Reid, Mohammad A. Mandegar

## Abstract

Recombinant adeno-associated virus (rAAV) is a widely used viral vector for gene therapy. Despite its clinical efficacy, the manufacturing of rAAV faces challenges in productivity and quality, leading to limited availability. To address the growing demand, next-generation process development should be informed by a mechanistic understanding of the cellular response to rAAV. In this study, we performed transcriptomic analysis of 5 cell lines with variable capacities for rAAV production. Using an intersectional approach, we assessed the transcriptional response to rAAV production and compared transcriptional profiles between high and baseline producers to identify possible targets for enhancing production. Modulation of cell cycle and nucleosome components suggested a reduction of proliferative capacity and a shift toward DNA replication to support rAAV production. During rAAV production, we observed upregulation of several core functions including transcription, stress response, and Golgi and endoplasmic reticulum organization. Conversely, inhibitors of DNA-binding proteins and mitochondrial components were consistently downregulated during rAAV production. We next performed a drug connectivity analysis of these results and identified 5 classes of drugs predicted to enhance rAAV production. Validation studies confirmed the efficacy of HDAC and microtubule inhibitors. Our data uncover novel and previously identified pathways that may enhance rAAV productivity, potentially enabling a path to engineer improved processes and cell lines for higher yields and better quality rAAV production.

**Graphical abstract:** 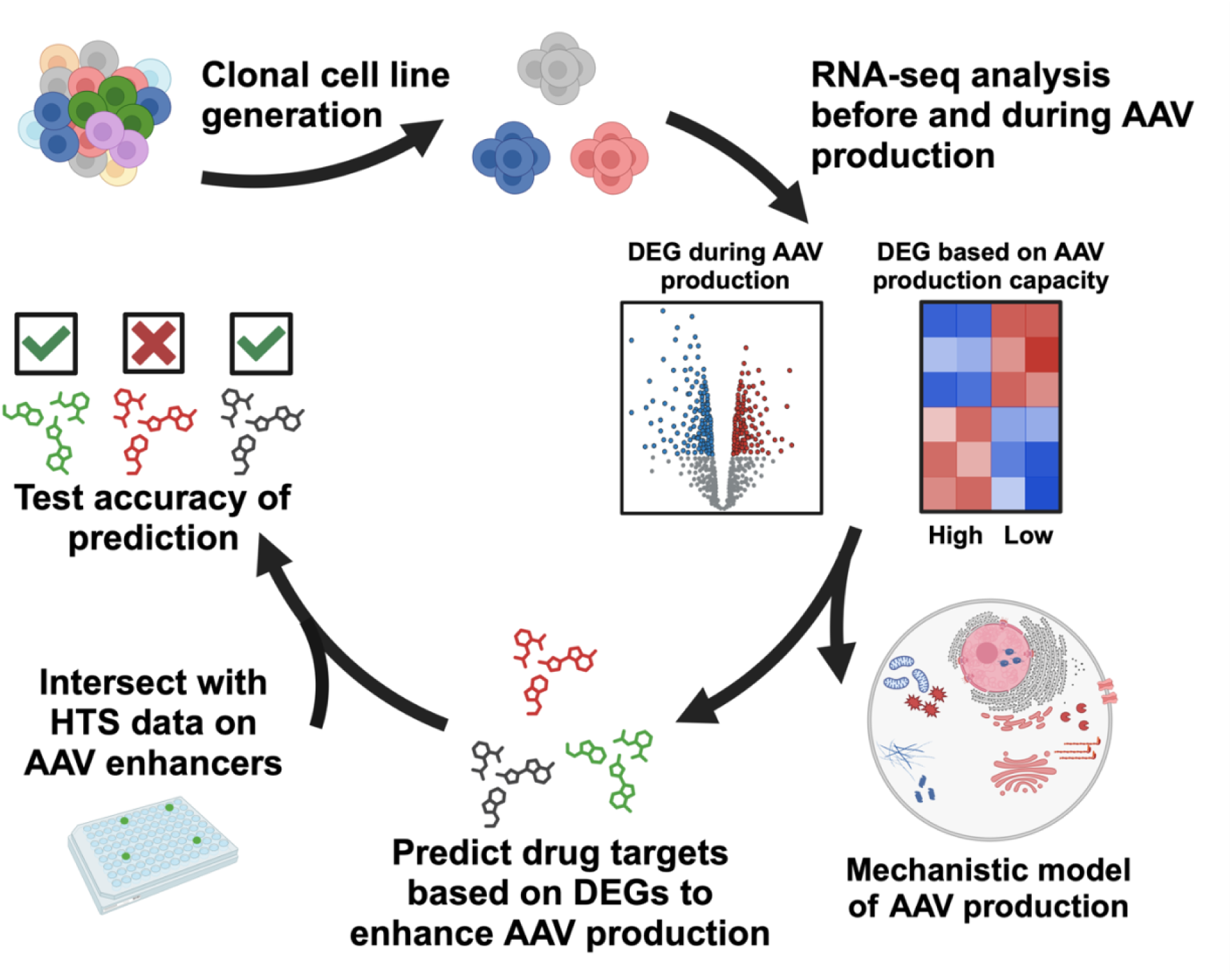

## Introduction

Adeno-associated viruses (AAVs) are non-pathogenic single-stranded DNA viruses approximately 4.7-kb long. Recombinant AAVs (rAAVs) have gained popularity in the gene therapy field due to their well-established safety profile, broad and tunable tropism and nonpathogenic nature (Pupo et al., 2022; Samulski and Muzyczka, 2014). As of 2024, we have seven commercially approved rAAV gene therapies (FDA, 2024), several programs in late stage clinical development, and hundreds in earlier stages of development (Kuzmin et al., 2021; Wang et al., 2024).

Given the promise of curative rAAV therapies, there is need for high doses of vectors especially for systemically delivered AAVs. Furthermore, there is growing emphasis on enhancing the quality of rAAV vectors through reducing plasmid- and host-derived impurities and partially packaged vectors (Chung et al., 2023; Wright, 2023). Widespread adoption of rAAV-based gene therapies has been hindered by challenges with production and manufacturing at scale.

Current methods of rAAV production result in relatively low volumetric yields, estimated to be 1000-4000-fold lower compared to monoclonal antibody production (Wright, 2023). Our limited biological understanding of cell-based rAAV production has led to inefficient manufacturing processes. rAAV molecules are complex, requiring careful assembly of 60 protein subunits to form the capsid, followed by incorporation of a single-stranded DNA molecule (Bennett et al., 2017; Onishi et al., 2023; Srivastava et al., 2021).

Given that most cell-based biomanufacturing platforms (Dumont et al., 2016; Tan et al., 2021) use cells outside of their evolutionary context to function as biologics production factories, it may not be possible to push rAAV production past physiological levels with currently off-the-shelf production platforms. The rAAV production process requires precise kinetics of expression and stochiometric ratio of replicase (Rep) and capsid (Cap) helper elements (Nguyen et al., 2021). Rep expression is known to induce cell cycle arrest (Berthet et al., 2005), and the helper gene E4 activates stress and DNA damage response, resulting to apoptosis (Hart et al., 2007). During viral replication and assembly, host cell defense and antiviral response pathways are activated (Chung et al., 2023), along with other unknown barriers that limit AAV production (Nguyen et al., 2021; Srivastava et al., 2021). By better understanding and eliminating these barriers, it may be possible to enhance the efficiency of viral replication and assembly.

To deepen our understanding of altered pathways during rAAV production, we analyzed RNA expression patterns using 5 different HEK293 cell lines in a time-series manner.

This revealed an array of cell line-independent changes in core pathways which may promote or inhibit AAV production when modulated. By applying pathway-drug connectivity analysis to our results, we identified potential pharmacological enhancers of rAAV production. Some targets have previously been implicated in rAAV production, while others were novel and unreported. Our study highlights the power of transcriptional profiling to identify genes and pathways as targets for cell and process engineering to improve rAAV production for clinical applications

## Results

### Clonal cell line generation with improved AAV9 yields

To ensure we identify robust and widely applicable transcriptional changes during AAV9 production, we used HEK293 cells from two separate sources. Expi293F (“293F” for short) were obtained from Thermo Fisher Scientific, and HEK293 cells (“AC1P” for short) from Cytion. We first developed standard AAV production protocols using triple plasmid transfection in both 293F and AC1P cells and monitored cell growth, cell viability and AAV production for 10 days after transfection (Figure 1 – figure supplement 1A-C).

**Figure 1.**
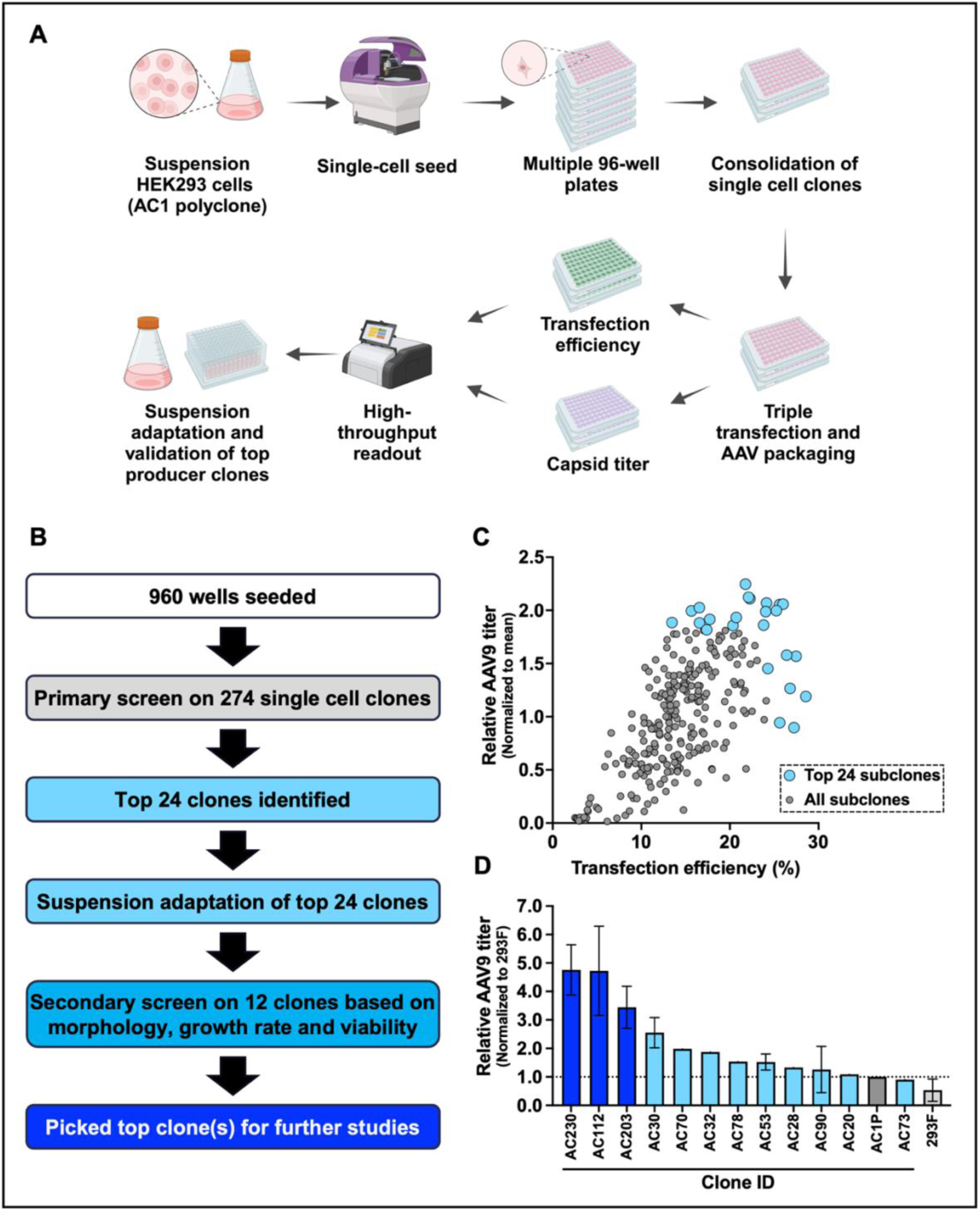
Clonal cell line generation with improved AAV9 yields. (**A**) Schematic of workflow for single-cell cloning and isolation of HEK293 clones with improved AAV production capacity. The Solentim VIPS™ single cell seeder was used to isolate clonal HEK293 cells using a two-step clonality verification process. (**B**) Subclone selection strategy for identification of top AAV9 producer clonal cells. (**C**) Approximately 300 single-cell derived HEK293 clones were identified based on growth, and top 24 performing colonies were selected based on transfection efficiency and AAV9 production. (**D**) After suspension adaptation, AAV productivity of the best suspension adapted clones was measured relative to AC1P (parental polyclonal cell line). n = 1–2 replicates per cell line. Data are shown as mean ± SD fold-change from AC1P (control).

We next isolated clonal cell lines from the AC1P (polyclonal) cell line using the Solentim VIPS™ (Verified in-situ plate seeding) single cell seeder to identify clones with higher AAV production potential (Figure 1A, B). During seeding and colony growth, a double imaging system (once at droplet dispense and subsequent whole well imaging) is used for clonality assurance (Figure 1 – figure supplement 1D). Approximately 300 clonally derived HEK293 colonies were screened for transfection efficiency and AAV9 production in adherent culture format (Figure 1C). We selected 24 top clones based on these parameters to convert to suspension format for further development and characterization. Cell viability and growth rate of these clones were measured during the suspension adaptation phase (Figure 1 – figure supplement 1E & F). After a 9-day suspension adaptation phase, the top 12 clones based on growth and viability were selected and expanded to 125 mL shake flask for secondary AAV9 production screens. Several of the clonal cell lines showed improved AAV9 production compared to 293F and the polyclonal AC1P parental line (Figure 1D).

### Cell line selection and RNA-Seq experimental design

We next selected the top 3 AAV producer clonal cell lines (AC230, AC112, AC203; Figure 1D) for transcriptomic analysis alongside 293F and AC1P. For each of these lines, we performed triple plasmid transfection in 125mL shake flask format and collected samples for RNA-seq analysis before transfection (0 hr) and during AAV9 production (24 and 72 hr after transfection) (Figure 2A). AAV9 vector genome (vg) titer was also measured from these samples at 72 hr (Figure 2B). Next, to identify genes associated with different AAV yield phenotypes, we classified cell lines into two categories: “base” producers (293F and AC1P) and “high” producers (AC112 and AC230). Base producers yielded an average of ∼2.4E9 vg/mL and high producers yielded an average of ∼1.1E10 vg/mL (Figure 2B).

**Figure 2.**
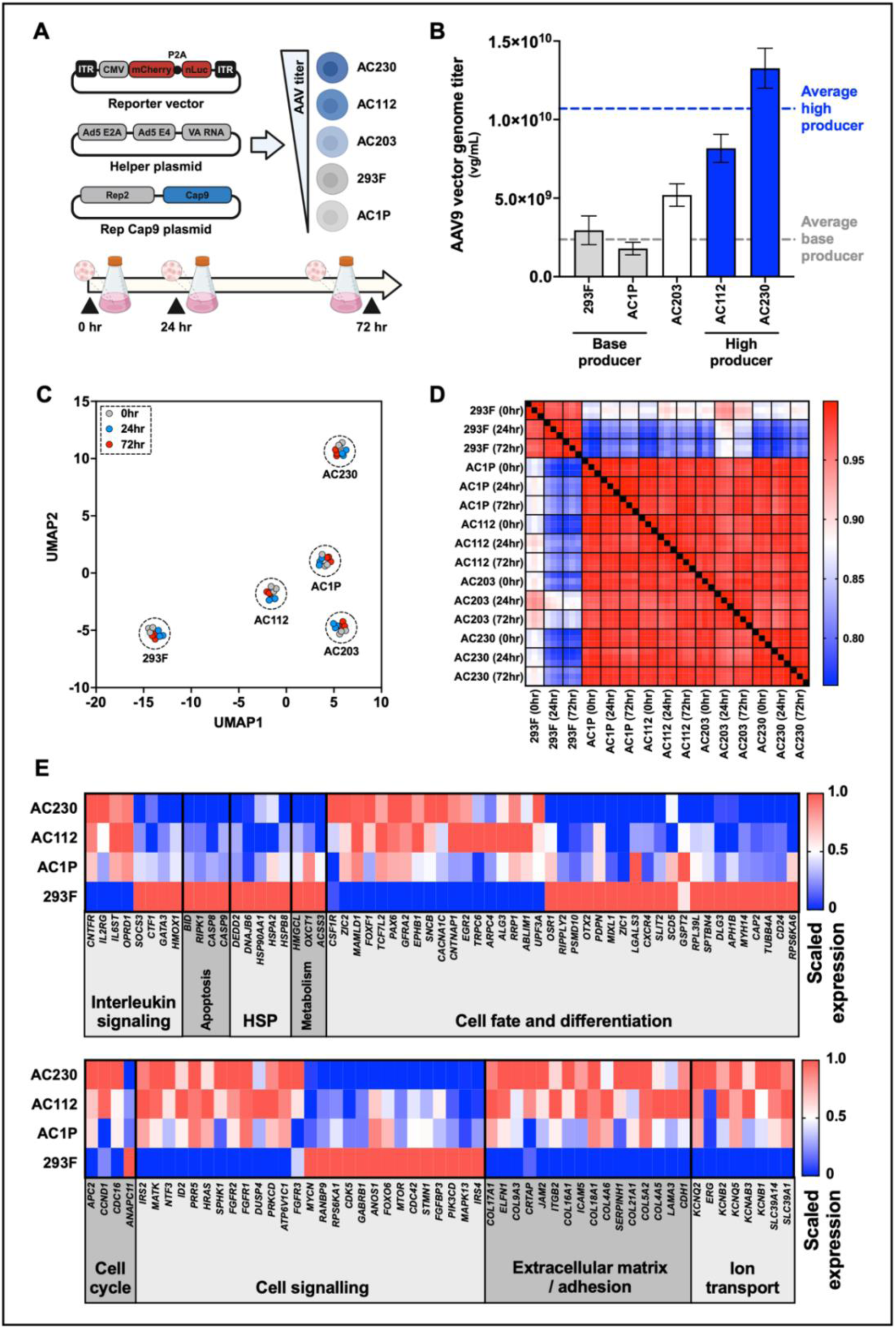
RNA-sequencing experimental strategy and top differentially expressed gene sets for high vs. baseline AAV producer cell lines. (**A**) Workflow and experimental design for RNA-sequencing analysis during AAV production. 293F, AC1P (polyclonal cell line) and 3 clonally derived cell lines from AC1P were used for the analysis. Samples were collected before transfection (0 hr) and at 24 and 72 hr after triple-plasmid transfection. 3 small-scale shake flasks were used per condition. (**B**) AAV9 vector genome titer was measured at 72 hr from the 5 cell lines. 293F and AC1P cells were designated as “base producers” with an average titer of ∼2.4E9 vg/mL. AC112 and AC230 were designated as high producers with an average titer of ∼1.1E10 vg/mL. n = 3 replicates per cell line. Data are shown as mean ± SD. (**C**) Uniform manifold approximation and projection (UMAP) and (**D**) gene correlation matrix of the 5 HEK293 cell lines show high reproducibility among replicates. RNA expression differences between cell lines are greater drivers of variability than differences between timepoints after transfection. (**E**) Heatmap of differentially enriched genes and functional categories in higher AAV producer cells (AC230 and AC122) compared to baseline producer cells (AC1P and 293F). Differentially enriched transcripts were manually categorized into cell cycle modulators, cell signaling, Extracellular matrix / adhesion, Ion transport, mitochondrial, heat shock protein, and cell fate and differentiation.

Bioinformatics workflow on raw data and quality control parameters are outlined in Figure 2 – figure supplement 1A–C. UMAP and RNA expression correlation analyses indicate high reproducibility among replicates and shows differences between cell lines are greater drivers of variability than differences between timepoints after transfection (Figure 2C, D). Additionally, AC1P transcriptional profiles are more correlated with its derived subclones than with the independently derived 293F line, as expected (Figure 2D and Figure 2 – figure supplement 1D).

### RNA-Seq analysis identifies differentially expressed pathways associated with improved AAV production

Our first aim was to identify gene expression patterns that are associated with improved AAV production. To this end, we compared gene expression patterns of the “high” producers (AC230, AC112) to the two “base” producers (AC1P, 293F) before transfection (0 hr) (Figure 2B). A total of 801 genes were significantly differentially expressed (|log_2_ (fold change) | > 0.5; adjusted P-value < 0.05) between base and high producers with 327 genes significantly upregulated in high producers, and 474 genes significantly downregulated in high producers (Supplementary file 1).

Next, we performed Reactome pathway analysis using the full list of differentially expressed genes to identify significantly enriched and depleted pathways in high versus base producers (Milacic et al., 2024). Using this analysis, we found that these genes commonly comprised pathways regulating cellular stress response (interleukin signaling, apoptosis, heat-shock), cell state (cell fate and differentiation, cell cycle), and extracellular matrix/adhesion organization and ion transport. We hypothesize that one or more of these pathways may contribute to enhanced AAV production in AC230 and AC112 (Figure 2E).

### AAV production is associated with transcriptional modulation of cell state, signal transduction, homeostasis, and extracellular interactors

Next, we focused our analysis on the subset of time-dependent expression differences which are shared across all 5 cell lines during AAV production. We identified significantly differentially expressed genes at 24 and 72 hr after transfection and compared to before transfection (0 hr). 499 genes were significantly differentially regulated at 24 hr, with 228 being upregulated and 271 downregulated. At 72 hr, 542 genes were differentially expressed, with 481 upregulated and 61 downregulated (Figure 2 – figure supplement 1E; Supplementary file 2).

Manual pathway analysis of differentially expressed genes revealed an array of fundamental host cell processes that were altered during AAV production, some of which overlap with processes described above when comparing gene expression between different AAV producer cell lines (Figure 2E). Notably, among the top upregulated genes during AAV production were those involved in cell fate and differentiation (eg. *CELSR3, TNFRSF12A, LIF*), signal transduction (eg. *INHBE, AMH, TNFRSF9*), cell cycle regulation (eg. *GADD45B, CEND1, NABP1*), protein homeostasis (eg. various heat-shock protein family members), and transcriptional activation (eg. *FOSB, ETV4, ETV5, RELB, MYB*). Among the most significantly downregulated factors were negative regulators of transcription (*ID1-3*) and extracellular matrix/cell adhesion molecules (eg. *HAS2, COL3A1*) (Figure 3A-D).

**Figure 3.**
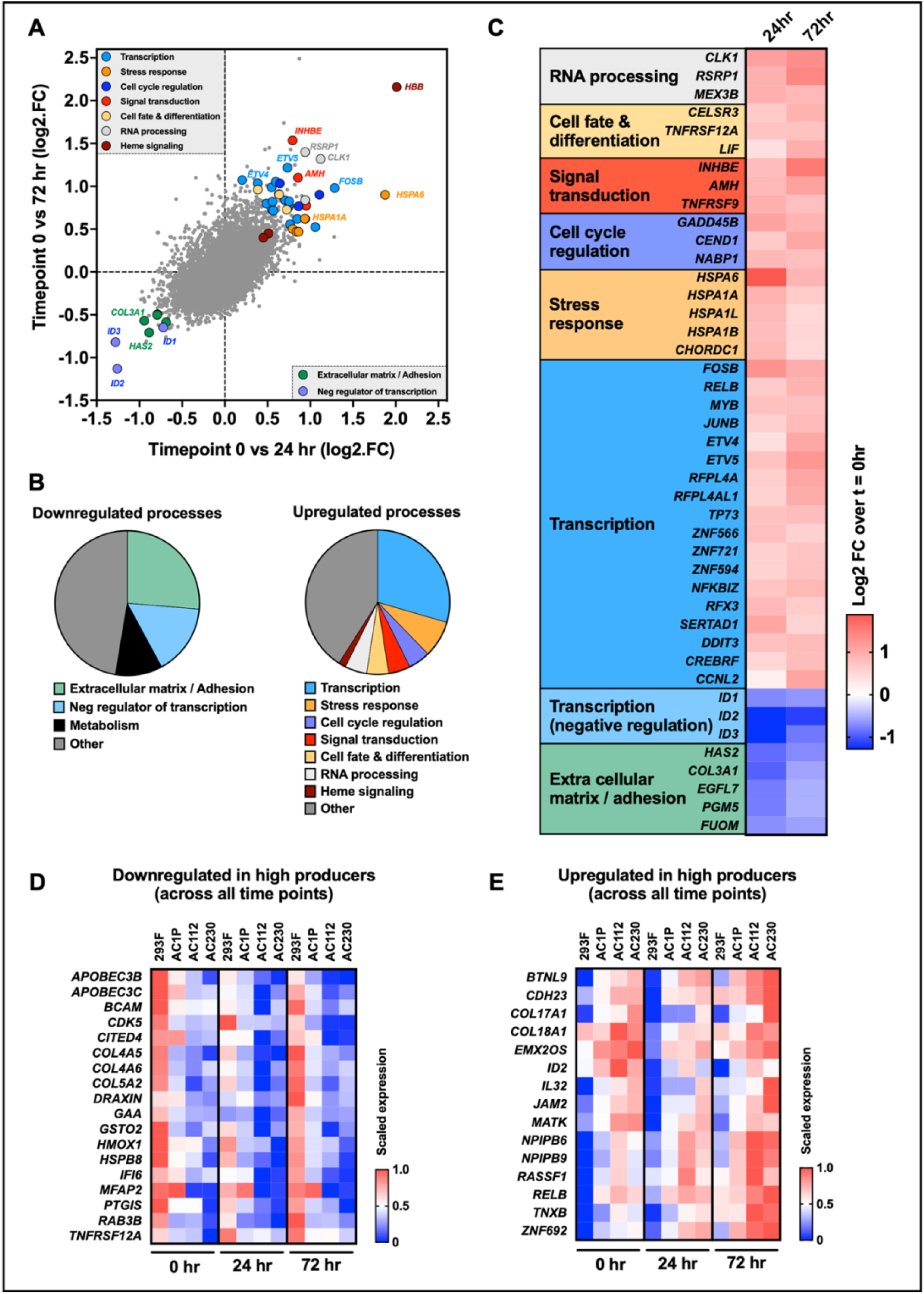
RNA-sequencing reveals modulation of transcription, protein homeostasis, signal transduction, and cell cycle regulation gene sets during AAV9 production. (**A**) Scatterplot comparing fold change gene expression at 72 hr vs. 24 hr during AAV production compared to 0 hr (before transfection). Data averaged from all 5 cell lines and 3 replicates per timepoint. Several gene families and genes of interest highlighted. (**B**) Pathway analysis of differentially expressed genes shows upregulated transcripts represent transcription factors, heat shock proteins, signal transduction molecules, cell cycle regulators, and Heme signaling molecules. Other upregulated processes include microtubule destabilizers, Golgi organization and vesicle transport. Downregulated transcripts include inhibitors of DNA-binding and extracellular matrix proteins. (**D**) Heatmap of most significant differentially expressed genes at 24 and 72 hr during AAV production. Data averaged from all 5 cell lines and 3 replicates per timepoint. Color scale represents log_2_ (fold change) expression from before transfection. (**E**) Heatmaps of results from intersection of genes modulated during AAV production with genes downregulated (left) and upregulated (right) in high AAV producer cell lines. Color scale represents log_2_ (counts per million), averaged across replicates and normalized from 0 to 1 for each gene.

Recent transcriptional studies have implicated many of the same pathways and processes we identified in AAV production, despite variations in study design including plasmid and cell sources, transfection conditions, and growth media (Supplementary file 3). Wang et al. (2023) identified several upregulated stress response and homeostatic proteins which align with our data, including *HBB*, *HSPA6*, and *CCL5*, as well as microtubule-interacting proteins including *STMN4* and *DNAH17* (Figure 3 – figure supplement 1A-C).

Another recent RNA sequencing-based study reported transcriptional upregulation of nucleosome assembly, inflammatory, viral response, and histone genes during AAV production (Chung et al., 2023). We observed similar trends in our dataset, with significant elevation of nucleosome assembly and histone genes and, with a few exceptions, a mild upregulation of inflammatory response and virus defense genes at 72 hr (Figure 3 – figure supplement 2A-C). Activation of only a handful of upstream regulators of differentially expressed genes reported by Chung et al. (2023) was detected in our study (Figure 3 – figure supplement 2D).

### Intersectional analysis reinforces transcription, cell proliferation, and extracellular matrix organization as key processes involved in AAV production

Ultimately, we aimed to isolate the genes which were both involved in AAV production and associated with higher AAV production. To this end, we next carried out an intersectional analysis of the genes which were modulated during AAV production and the differentially expressed genes we identified between our high vs. base AAV producers. For this analysis, we first identified all unique genes which were either up- or downregulated during AAV production between any two timepoints. Second, we identified genes which were either upregulated or downregulated at all timepoints (before transfection, 24 and 72 hr after transfection) in high AAV producing cell lines.

This produced a list of 1143 unique genes which were modulated during AAV production, 110 genes which were upregulated in high producers at all timepoints, and 245 genes which were downregulated in high producers at all timepoints (Supplementary file 4). Finally, we intersected the 1143 AAV production-related genes with the 110 upregulated- and 245 downregulated-in-high-producer genes. This enabled us to identify a subset of targets which we hypothesize to enhance AAV production in HEK293 cells when artificially perturbed (Figure 3E).

Among those genes which were relevant to AAV production and upregulated in high producers were negative regulators of proliferation (*BTNL9, EMX2O9, MATK, RASSF1*), transcription factors (*RELB*, *ZNF692*) and nuclear pore complex components (*NPIPB6, NPIPB9*). Genes which were both downregulated in high producers and differentially regulated during AAV production included viral DNA-targeting cytosine deaminases (*APOBEC3B/C*), positive regulators of proliferation (*CDK5, MFAP2, PTGIS, RAB3B, TNFRSF12A*) and stress-responsive genes (*GSTO2*, *HMOX1, HSPB8, IFI6*). A variety of cell adhesion and extracellular matrix molecules which were modulated during AAV production exhibited either upregulation (*CDH23, COL17A1, COL18A1, JAM2, TNXB*) or downregulation (*BCAM, COL4A5, COL4A6, COL5A2*) in high AAV producer cells.

### A model for AAV production based on coordinated regulation of multiple cellular processes

To gain a deeper understanding of the regulatory events underlying AAV production, we identified several significantly up- and downregulated cellular processes based on a gene ontology (GO) analysis. Glutathione-related processes were among the most significantly downregulated GO terms, suggesting a role for oxidative stress in AAV production. Mitochondrial processes, particularly electron transport, NADH dehydrogenase, and respiratory chain complex I, were also significantly downregulated. This was accompanied by a downregulation among genes involved in collagen fibril organization, again implicating extracellular interactions in AAV production. Conversely, Amino acid transport, stress-responsive RNA polymerase activity, endoplasmic reticulum stress, and Golgi cisterna membrane components were among the most significantly upregulated GO terms during AAV production (Figure 4A, B). We also observed a concordant upregulation of nuclear pore complex and histone components, reflecting the importance of nuclear transport and viral DNA replication during AAV production (Figure 4B) (Junod et al., 2021; Nicolson and Samulski, 2014).

**Figure 4.**
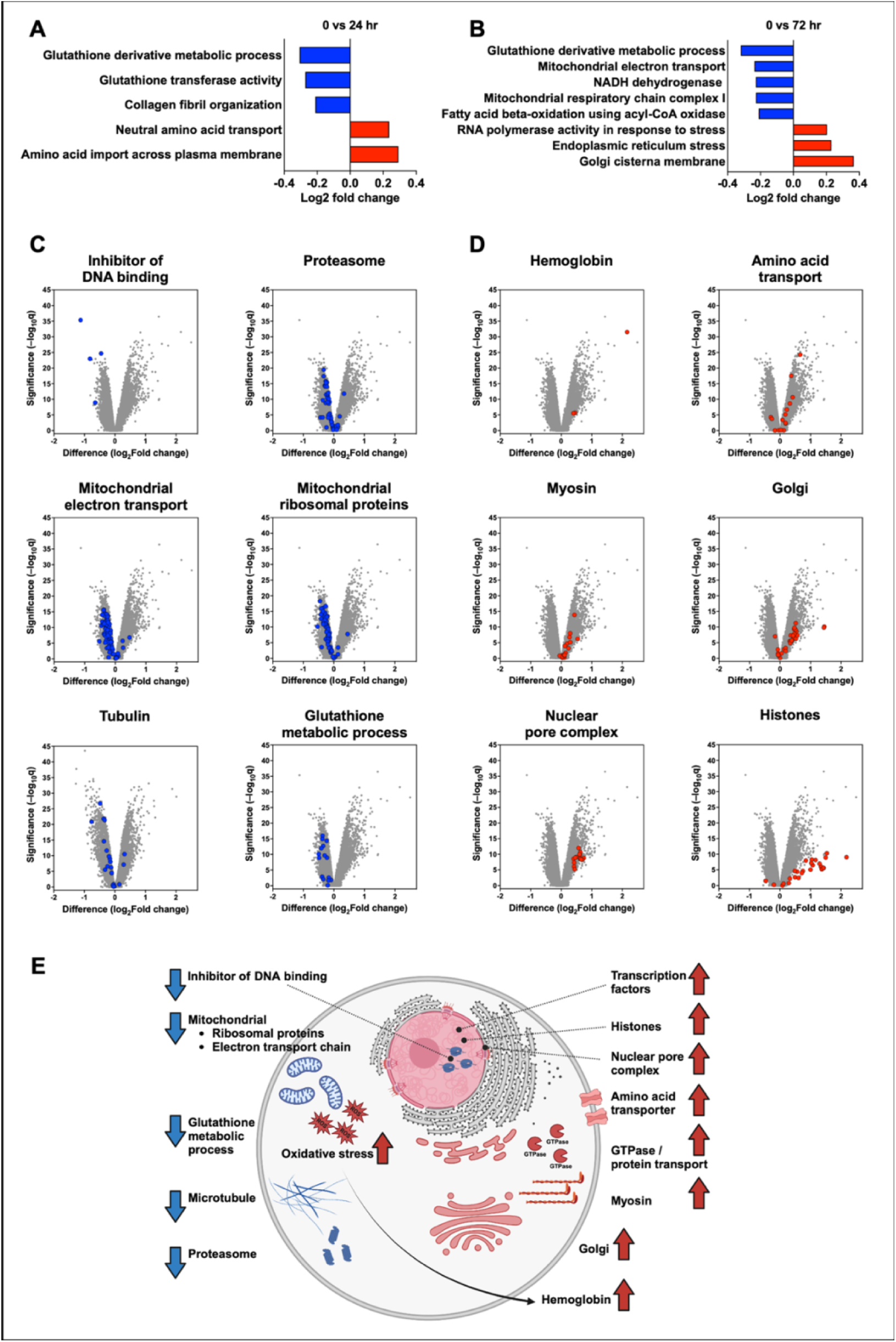
Selection of differentially expressed cellular components and processes incorporated into a model for AAV production. (**A, B**) Gene ontology (GO) term analysis for the 24 hr (**A**) and 72 hr (**B**) timepoints reveals downregulation among mitochondrial, collagen, and glutathione and fatty acid metabolic processes during AAV production. Significantly upregulated GO categories include amino acid transport, stress response, and Golgi cisterna components. (**C**) Volcano plots of select processes reveal coordinated downregulation among inhibitor of DNA binding, proteasome, mitochondrial electron transport, mitochondrial ribosomal proteins, tubulin, and glutathione metabolic process genes during AAV production. (**D**) Volcano plots of select processes reveal coordinated upregulation among hemoglobin, amino acid transport, myosin, Golgi, nuclear pore complex, and nucleosome assembly genes during AAV production. (**E**) Model of activated and repressed pathways during AAV production points to targets that could be modulated to enhance AAV yield and quality.

We next filtered our list of differentially expressed genes at 72 hr after transfection for gene sets within each of these GO categories. In alignment with our earlier observation of upregulation among transcriptional activators (Figure 3C & D), we observed strong and coordinated downregulation of inhibitors of DNA binding genes (*ID1*-*ID3*). Other negative regulators of transcription which were downregulated by 72 hr included *MAFB* and *SMAD6* (Supplementary file 2). Additionally, most molecules compromising proteasome (e.g. *PSMD9*, *PSMA6*) and mitochondrial ribosomal proteins and electron transport chain were coordinately downregulated, suggesting broad downregulation of protein turnover and energy production pathways during AAV production (Figure 4C & Figure 4 – figure supplement 1A). This observation, paired with overall upregulation among amino acid transport (e.g. *SLC3A2*, *SLC1A3*), Golgi (e.g. *GOLGA8A*, *GOLGA8B*), nuclear pore complex (e.g. *NPIPA1*, *NPIPB9*) and nucleosome assembly genes (e.g. *H1-2*, *H2BC5*) (Figure 4D & Figure 4 – figure supplement 1B) highlights a cellular transition away from oxidative energy production and toward biomolecular synthesis essential for AAV production. Interestingly, we observed opposing regulation of two cytoskeletal components: tubulin, which was generally downregulated, and myosin, which was generally upregulated. This highlights a central role of specific cytoskeletal rearrangements for intracellular transport of viral particles during AAV production.

Based on these results, we have constructed a simple model of activated and repressed pathways during AAV production (Figure 4E). This model proposes specific cellular processes which may be targeted for perturbation with the goal of enhancing AAV production for therapeutic use.

### Drug connectivity analysis points to cell cycle and epigenetic modulators as enhancers of AAV9 production

Next, we leveraged our RNA-seq data to identify targets that could be pharmacologically modulated to enhance AAV production. Our workflow and triaging strategy to identify putative pharmacological enhancers of AAV production is outlined in Figure 5 – figure supplement 1A. First, we identified enriched and depleted targets or mechanisms of action during AAV production. Approximately 1000 compounds were predicted to modulate the enriched pathways. We next filtered this primary list to only include annotated drugs with known (or putative biological targets) with a *P* <0.01 and a normalized enrichment scores (NES) greater than 1.3 or less than **–**1.3. Examples of enriched targets and pathways included protein synthesis and tubulin polymerization, while depleted pathways included histone deacetylases (HDAC) and cell cycle and cell signaling kinases (Figure 5A, B, Figure 5 – figure supplement 1B, C).

**Figure 5.**
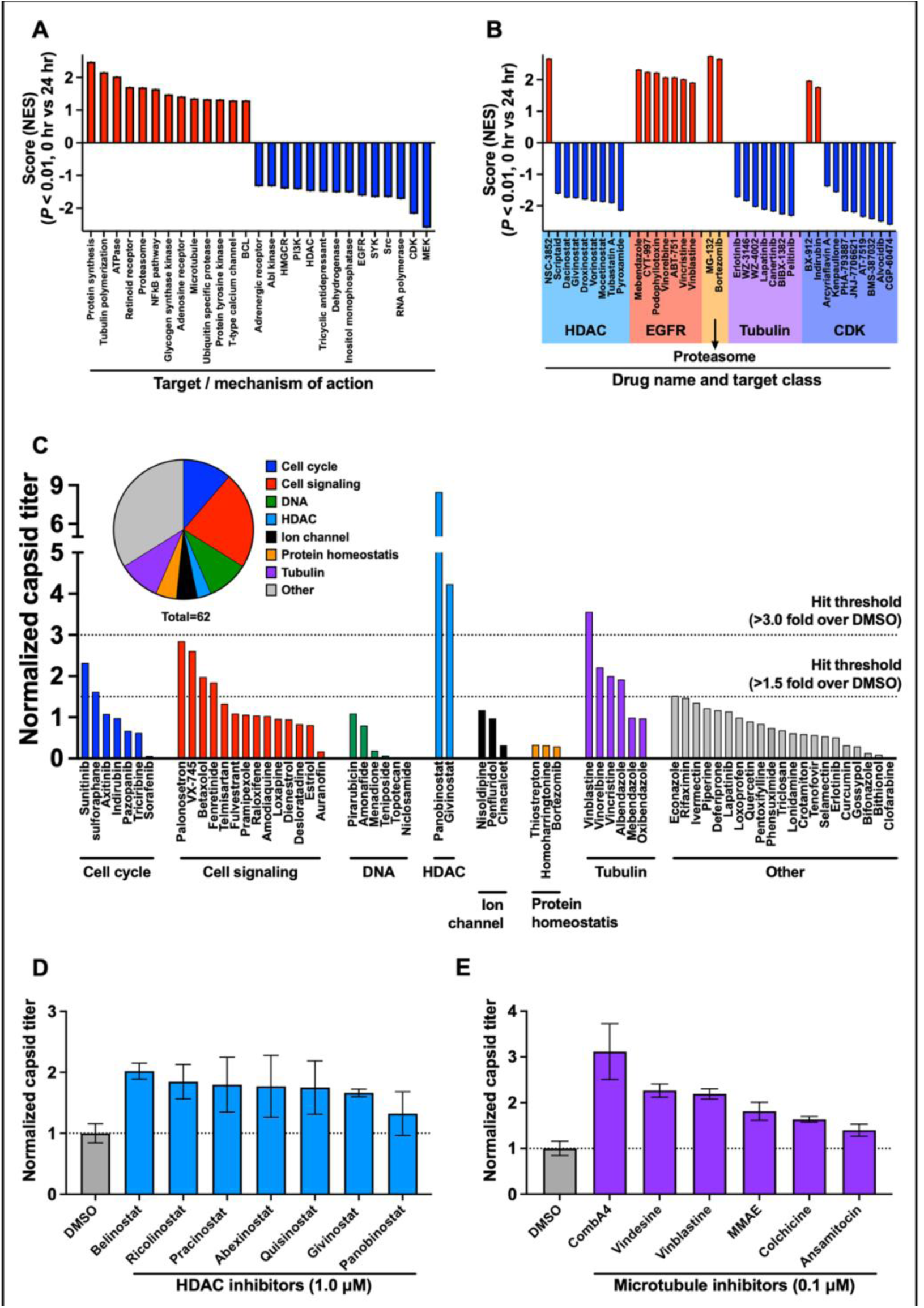
Pathway analysis identifies druggable pathways that may enhance AAV production when modulated. (**A**) Normalized enrichment score (NES) with P-values <0.01 was used to identify enriched and depleted targets or mechanisms of action during the early stages of AAV production (0 vs 24 hr). Examples of enriched pathways included protein synthesis and tubulin polymerization, while depleted pathways included histone deacetylases (HDAC) and various classes of kinases (mostly involved in cell cycle and cell signaling). (**B**) Normalized enrichment score (NES) with P-values <0.01 of drugs with known targets affecting enriched and depleted pathways during AAV production (0 vs 24 hr). Enriched drug classes include HDAC, EGFR, proteasome, tubulin and cyclin dependent kinase modulators. (**C**) Data from 62 drugs which were tested at 1µM dose and screened for fold increase AAV9 capsid titer compared to the DMSO controls. From the 62 drugs, 12 increased AAV9 production 1.5-fold above baseline (∼19% hit rate), while 3 out of 62 increased AAV9 production 3-fold above baseline (∼5% hit rate). (**D, E**) Validation studies were carried out using 7 HDAC (**D**) and 6 microtubule (**E**) inhibitors applied at the time of transfection. These led to a 1.5- to 3-fold increased AAV yield compared with DMSO control. n = 2–6 replicates per treatment. Data are shown as mean ± SD fold-change from DMSO control.

We next identified annotated compounds that modulate these pathways. A total of 266 annotated compounds were predicted to influence these pathways during AAV production. Out of the list of triaged compounds, 62 were available in our small molecule screening library (Grafton et al., 2023). These compounds target targets and cellular processes including cell cycle, cell signaling, DNA replication and synthesis, histone deacetylase (HDAC), ion channels, protein homeostasis, and tubulin. We tested these 62 compounds at 1 µM dose and screened for AAV9 capsid titer (Figure 5C).

From the test set of drugs tested, 12 increased AAV9 production 1.5-fold above baseline (∼19% hit rate), and 3 out of 62 increased AAV9 production 3-fold above baseline (∼5% hit rate). We further validated our results using top 2 mostly well-defined target classes: HDAC and microtubule. Validation studies were carried out using 7 HDAC and 6 microtubule inhibitors applied at the time of transfection. These led to up to ∼2 and ∼3-fold increase in AAV yields compared with DMSO control (Figure 5D, E).

## Discussion

In this study, we combined transcriptional analysis with pharmacological perturbations to improve our mechanistic understanding of key pathways during AAV production. We incorporated 5 HEK293 cell lines with varying capacities for AAV production and used RNA-sequencing to identify genes and pathways that are both modulated during AAV production, and differentially expressed between base and high AAV producers. Using this intersectional strategy, we identified a subset of differentially expressed host cell molecules which we hypothesize may enhance AAV yield when ectopically expressed or depleted.

Among the core cellular functions upregulated during AAV production were transcriptional activation, cellular stress response, signal transduction, and cell cycle regulation. On the other hand, we found that mitochondrial ribosomal and electron transport components, proteasomal proteins, glutathione metabolic process modulators, and transcriptional repressors were generally downregulated during AAV production. We found that many of these processes were altered when comparing expression profiles between base and high AAV-producing cell lines. Using these results, we developed a general cellular model for AAV production characterized by a complex shift in cell state, with widespread modulation of energy production and biosynthesis.

With the knowledge and key pathways identified in our study, we used drug-pathway interaction analysis to predict, triage, and subsequently test the effects of pharmacological modulators of altered pathways during AAV production. A subset of these predicted compounds were tested, and up to 19% of the tested compounds improved AAV production 1.5-fold above baseline. This is a substantially higher hit rate than if we had performed an unbiased high-throughput screen where typical hit rates range from 0.1 – 1% (Barnes et al., 2021; Dreiman et al., 2021; Yang et al., 2022). Our approach demonstrates that selective screening of compounds using a transcriptomics-informed strategy results a higher success rate than a hypothesis-free approach in increasing viral production. We further exemplified our approach by identifying and subsequently validating HDAC and microtubule inhibitors as enhancers of AAV production. Our findings demonstrate that targeting epigenetics, activation of transcriptional machinery and cell cycle modulation are important mechanisms in promoting AAV production.

AAV production via plasmid transfection is associated with a broad range of transcriptional responses in the host cell. Modulation of transcription resulting directly from transfection is generally short-lived and returns to baseline by 24 hr after transfection (Chung et al., 2023). During AAV production, host cell transcriptional machinery must accommodate the production of both viral proteins and host cell proteins involved in downstream homeostatic responses. This was evident in our data, where we observed upregulation of many transcription factors during the mid-late phase of AAV production. These include *ETV4* and *ETV5*, which bind to the enhancer of adenovirus E1A gene, and act as transcriptional activators (Higashino et al., 1993). We also found upregulation of *FOSB, RELB, MYB* DNA-binding factors that are known proto-oncogenes and control proliferation and cell differentiation (Lipsick and Baluda, 1986; Nakabeppu and Nathans, 1991; Ryseck et al., 1992).

Transcriptional upregulation of histones has also been reported during AAV production (Chung et al., 2023) further indicating that epigenetic modulation of transcriptional machinery is a key mechanism in enhancing viral replication and production. Previous studies in addition to ours have shown that HDAC inhibitors can increase AAV (Scarrott et al., 2023; Wada, 2022) and lentivirus production (Paszkiet et al., 2022). This is likely due to their capacity to stimulate viral gene transcription through chromatin remodeling (Gallinari et al., 2007).

Furthermore, we observed a coordinated downregulation of negative regulators of transcription during AAV production, particularly inhibitor of DNA binding proteins *ID1*-*3*, *MAFB,* and *SMAD6* which may be acting as barriers to viral replication. *ID* isoforms are also thought to play a role in cell growth, senescence, and differentiation (Lasorella et al., 2014). MAFB represses erythroid genes including the transferrin receptor, which is essential for heme synthesis (Sieweke et al., 1997). This is in alignment with our observed upregulation of *HBB* during AAV production. SMAD6 acts as a transcriptional corepressor to inhibit BMP signaling in the nucleus via a negative feedback loop (Bai et al., 2000; Imamura et al., 1997). It is possible that *SMAD6* downregulation is a compensatory response to the reduction in *ID1-3,* which are downstream effectors of BMP signaling (Peng et al., 2004; Yang et al., 2013).

We observed opposing regulation of proliferative genes which were involved in AAV production, with a general upregulation of inhibitors of proliferation, and downregulation of enhancers of proliferation among high AAV producers. This is reflective of the cell cycle dependence of AAV production, with AAV most effectively produced during the S/G2 phase (Franzoso et al., 2017). This, coupled with a coordinated upregulation of histones during AAV production (also previously reported by Chung et al. (2023)), is characteristic of the S phase of the cell cycle when viral DNA replication is most efficient (DeLisle et al., 1983; Polo and Almouzni, 2005). Tubulin and myosin are also involved in mitotic spindle formation, thus affecting cell cycle progression and proliferation (Ferreira et al., 2018; Shahid-Fuente and Toseland, 2023). We observed opposing regulation of these families during AAV production, with tubulin isoforms generally downregulated and myosin isoforms upregulated. It is possible that in addition to modulating the cell cycle, myosin is also used during AAV production to transport viral cargo during production.

Modulating cell cycle, particularly arresting cells in the G2/M phase using microtubule inhibitors such as Nocodazole (Scarrott et al., 2023), has been shown to boost AAV titers. A gain-of-function screen using both CRISPR activation and transgene overexpression showed spindle and kinetochore associated complex subunit 2 (SKA2), and inositol 1, 4, 5-trisphosphate receptor interacting protein (ITPRIP) increased AAV titer by altering the host cell cycle to increase AAV vector genome replication (Barnes et al., 2021). Cell cycle synchronization induced by temperature shift has also been shown to improve AAV (Coplan et al., 2024) and adenovirus (Ferreira et al., 2009) production. A slower cell growth rate may be beneficial during AAV production, as it reduces the rate of plasmid loss and dilution (Nguyen et al., 2021). Furthermore, by slowing down the cell’s growth rate, less cellular machinery is diverted to processes such as host cell DNA replication and protein synthesis, thereby freeing cellular resources for vector amplification, replication, and capsid assembly.

AAV production in our cell lines was associated with widespread modulation of various signaling pathways, including kinases and other elements involved in growth factor signaling. Notably, mitogen-activated protein kinases (MAPKs) are critical signaling molecules that are known to regulate cell cycle, differentiation, and apoptosis (Wilkinson and Millar, 2000; Zhang and Liu, 2002). Negative regulation of MAPK activity has been reported during AAV production (Chung et al., 2023). In addition, dysregulation of MAKP signaling using miR-431 and miR-636 has been reported to improve AAV2 assembly and potency by > 3-fold (Khan et al., 2020). These findings underscore the intricate interplay between cell cycle regulation and signaling pathways during AAV production and may offer strategies to enhance AAV vector yields.

During viral production, cells undergo various forms of cellular stress which evoke an array of responses via heat shock, unfolded protein response (UPR), oxidative stress, and immune response pathways (Balakrishnan et al., 2013; Chung et al., 2023; Foo et al., 2022; Wang et al., 2023; Zhang and Yu, 2022). Based on our data, some of the most significantly upregulated transcripts during AAV production were heat shock proteins (HSPs), including HSPA6, HSPA1A, HSPA1L, HSPA1B, and the HSP90 co-chaperone CHORDC1. HSPs have been shown by others to be activated by viral infection and may act to suppress viral replication, or they may be modulated by the virus itself to promote its replication (Gano and Simon, 2010; Zhang and Yu, 2022). It is possible that HSP expression is upregulated during AAV replication in part due to a large influx of immature proteins into the ER and Golgi complex, thus evoking a UPR. This is evidenced in our dataset by modulation of transcription factors which affect UPR, including ATF6B and several CREB-related factors (Supplementary file 2) (Adachi et al., 2008; Zhang et al., 2013).

Oxidate stress, due to mitochondrial damage (Foo et al., 2022), is also known to cause upregulation of hemoglobin (Liu et al., 2011). In alignment with this, we observed upregulation of *HBB*, *HBA1* and *HBA2* at 24 and 72 hr after transfection (Figure 4 - Supplementary figure 1). Conversely, heme oxygenase 1 (*HMOX1*) was significantly downregulated during AAV production and in our high AAV-producing cell lines. *HMOX1* is a stress response gene, and its product catalyzes the degradation of heme, an important cofactor of hemoglobin and various enzymes (Schweitzer-Stenner, 2022). The byproducts of heme degradation include reactive oxygen species scavengers which exhibit anti-inflammatory effects and can inhibit viral replication (Espinoza et al., 2017).

Activation of innate immune response and apoptotic pathways via interferons has been reported during AAV production (Chung et al., 2023; Wang et al., 2023). Downstream inhibition of interferon signaling using Ruxolitinib (JAK1-TYK2 inhibitor) blocks JAK-STAT signaling, and has shown to double AAV yields in low-producing cell lines (Kahlig et al., 2024). A separate report revealed that double knockout of two key regulators of apoptosis (*BAX* and *STAT1)* in HEK293 cells increased AAV5 titers by 1.8-fold (Turner-Gillies et al., 2024). In line with these findings, our high AAV-producing cell lines exhibited downregulation of genes encoding other key apoptotic factors including *BID, CASP8,* and *CASP9,* potentially enabling these cells to survive under enhanced stress associated with AAV production (Galluzzi et al., 2018). These results suggest inhibition of stress response pathways and apoptosis may be beneficial in promoting viral yields.

In conclusion, our study elucidates the complex transcriptional and cellular responses involved during AAV production. Through integration of pathway analysis and pharmacological screening, we show that targeted modulation of specific pathways such as transcriptional regulation and cell cycle control can enhance AAV production.

While our study advances our mechanistic understanding of AAV production and highlights potential strategies for enhancing AAV production, our study is limited in at least the following aspects.

Most commercially available bioactive compound libraries are enriched in anti-cancer drugs that either directly or indirectly affect cell cycle. Hence the scope of our search space is limited by the diversity of the chemical space, which does not probe all potential influential pathways and targets. Use of a fixed compound concentration at single dose (1 μM) in our initial screen limited our ability to identify potentially efficacious compounds that operate above or below this concentration. In addition, variability in gene expression patterns among different HEK293 cell lines suggest that our findings may not be universally applicable to all cellular production systems. This is especially relevant when using different plasmid systems, media formations, feed strategies, and transfection reagents.

Hence, further research is needed to expand the range of targetable pathways by optimizing compound dosage and implementing genetic screens via CRISPR/Cas9, siRNA, and transgene overexpression. These steps will be crucial in translating our hypotheses into practice and effectively scaling up from a laboratory setting to large-scale bioreactor systems. By addressing these challenges, we can enhance the reliability and efficiency of AAV production, paving the way for enhanced AAV production processes.

## Materials and Methods

### Adherent cell culture and suspension adaptation of HEK293 cells

Adherent polyclonal HEK293 cells (“AC1P” for short) (Cytion) were expanded in DMEM, high glucose, GlutaMAX™ Supplement, pyruvate (Thermo Fisher Scientific) supplemented with 10% fetal bovine serum (FBS) (Thermo Fisher Scientific) in a 37°C, 5% CO_2_ incubator. Once cells had grown to ∼80% confluency, the media was carefully aspirated and replaced with Expi293™ expression medium (Thermo Fisher Scientific) and incubated for 48 hr in the adherent format at 37°C, 5% CO_2_. After 48 hr of cell adaptation to Expi293™ expression media, cells were resuspended in 2 mL fresh media and placed on an orbital shaker (125 rpm, 19-mm orbital throw) in a 37°C, 5% CO_2_ incubator. After 2 days of suspension adaptation, cell viability and density were measured using a Countess 3 and Trypan Blue (Thermo Fisher Scientific) and cells were maintained at 0.5–5 M/mL. All cell lines tested negative for mycoplasma using a MycoAlert Mycoplasma Detection Kit (Lonza).

### Maintenance HEK293 cells in suspension prior to transfection

Expi293F^TM^ HEK cells (“293F” for short) (Thermo Fisher Scientific) and HEK293 (“AC1P” for short) (Cytion) were maintained in suspension in Expi293™ expression medium on an orbital shaker (125 rpm,19 mm orbital throw) in a 37°C, 5% CO_2_ incubator. Cell growth and viability was monitored for 7-10 days until cells had reached a density between 2.5 – 5 M/mL and >95% viability prior to transfection. Cells were then centrifuged at 500 g for 5 minutes. Spent media was aspirated and the cell pellet was resuspended in the appropriate volume of Expi293™ expression medium to achieve a density of 2.5 M/mL in a 30 mL shake flask.

### Clonal cell line generation and screening

To generate and screen clonal cell lines for enhanced AAV production, we resuspended AC1P cells to 9,500 cells/mL in DMEM, high glucose, GlutaMAX™ Supplement, pyruvate (without FBS) after ensuring >90% viability. We carried out single cell seeding into Corning Costar 96-well flat-bottom cell culture-treated plates (cat. number 3596) using the Solentim VIPS apparatus according to recommended instructions. Each seeded well was imaged immediately upon seeding before filling the well with DMEM+10% FBS. After seeding all wells, they were imaged again using the “verify clonality” function, and this was repeated each day for 5 days. Starting on day 5, we imaged wells using the “monitor colony growth” function and repeated this every other day until about 2 weeks after seeding.

Once colonies were 65-80% confluent, we consolidated them into new 96-well plates if they exhibited clear evidence of clonality (a single colony derived from a single cell at time of seeding). We then distributed consolidation plates among 3 sister plates for expansion, transfection, and cryopreservation. Transfection plates were pre-treated with fibronectin (1:1000 in PBS; STEMCELL Technologies) prior to transferring clonal colonies.

### AAV harvest and purification

The triple transfection system uses a pHelper plasmid, a RepCap9 plasmid and an ITR containing reporter plasmid. The pHelper plasmid contains Ad5 E4 and Ad5 E2A and Ad5 VA-RNA. The RepCap9 plasmid expresses AAV2-derived Rep78 and Rep68 under the control of the P5 promoter, Rep52 and Rep40 under the control of the P19 promoter and VP1, VP2 and VP3 derived from AAV9 under the control of the P40 promoter. The ITR-containing reporter plasmid includes mCherry-P2A-nanoluciferase (NLuc) driven by a CMV promoter. After transfecting cells using TransIT-VirusGEN (Mirus Bio), we allowed cells to produce AAV for up to 72 hr before harvesting samples for titer analysis. Cells were lysed via three rounds of freezing on dry ice followed by thawing at 37°C. After mixing lysed samples, cell debris was removed by centrifuging at 1000g for 10 min, and AAV-containing supernatant was saved for processing and stored at -80°C.

### Determination of capsid titer using ELISA

A Streptavidin coated high-capacity plate (Thermo Fisher Scientific) was coated for 1 hr at room temperature with the CaptureSelect™ Biotin anti-AAV9 conjugate (Thermo Fisher Scientific), at a dilution of 1:10,000 in PBST. After incubation, the capture antibody solution was aspirated from each well of the plate, and each well was washed times with 150 µL of PBST. 100 μL of either standard or sample was added to the plate. A 5–7-point standard was used in all studies using the AAV9 empty capsid standard (Progen). The plate was covered with foil sealing film and incubated on a shaker for 1 hr at room temperature. After incubation, the entire sample was aspirated off, and the plate was washed 3 times with 150 µL of PBST. CaptureSelect™ HRP anti-AAV9 conjugate (Thermo Fisher Scientific) was diluted 1:50,000 in PBST and 100 µL of the diluted detection antibody was added to each well. The plate was covered with foil sealing film and incubated on a shaker for 1 hr at room temperature. After incubation, the HRP anti-AAV9 conjugate solution was aspirated and washed 3 times with 150 µL of PBST. 100 µL of TMB ELISA substrate (highest sensitivity) (Abcam) was added per well and incubated for 2-5 minutes before adding 100 µL the ELISA stop solution (Thermo Fisher Scientific). Absorbance was measured on the Varioskan microplate reader at 450 nm (Thermo Fisher Scientific).

### Determination of vector genome titer using qPCR

Purified or crude viral supernatant was DNase I treated for 90 minutes at 37°C according to manufacturer’s instructions (New England BioLabs). qPCR was performed using a custom FAM probe against Nano Luciferase (Thermo Fisher Scientific) and TaqMan fast advanced master mix (Thermo Fisher Scientific) on a QuantStudio 6 Flex real-time PCR system (Thermo Fisher Scientific). Plasmid DNA was used from a concentration of 1 ng/µL to 0.0001 ng/µL to generate a standard curve.

### RNA preparation and sequencing

For each cell line and replicate, 2.5×10^6^ cells were removed before transfection (0 hr), as well as 24 and 72 hr after transfection. Cells were pelleted, flash frozen, and stored at -80°C. Total RNA was extracted using Direct-zol-96 RNA Kit (Zymo Research) according to the manufacturer’s instructions. RNA concentration was quantified using a NanoDrop spectrophotometer (Thermo Fisher Scientific) and RNA quality was assessed via High Sensitivity RNA Screen Tape Analysis (Agilent, performed via SeqMatic). Illumina stranded mRNA library preparation and sequencing (NovaSeq V1.5, S1, 100 cycles) were performed by SeqMatic. On average, 42,271,540 ± 7,224,160 reads (100-base pair, single-end) were generated per sample. We deposited our RNA-seq data on the Gene Expression Omnibus (GEO) database: GEO Submission (GSE269485).

### Differential expression analysis

We performed quality assessment on sequence reads using FastQC (Andrews, 2010) and aligned reads using Hisat2 (Kim et al., 2019) to the human genome (GRCh38.110). After generating genomic alignments, we used Subread’s featureCounts function to generate gene-level read counts (Liao et al., 2014). On average, Hisat2 aligned 95.6 ± 1.9% of reads to the genome one or more times, and 97.8 ± 1.2% of these reads were successfully assigned to genomic features.

After obtaining high-quality sequence alignments and counts, we generated an expression matrix which was submitted to the BigOmics platform for differential expression analysis (Akhmedov et al., 2020). Raw counts were converted into counts per million (CPM) and log_2_ normalized for downstream analysis. Protein-coding genes with at least 1 CPM across two or more samples were included in the analysis. By default, the BigOmics platform uses a combination of statistical tests for differential expression via DESeq2 (Wald), edgeR (QLF), and limma (trend).

Initial t-SNE clustering and differential expression analyses were performed within the BigOmics web environment and results were exported for visualization. Pairwise gene expression correlations between samples were calculated in R using the Pearson method. Multiple contrasts were generated in the BigOmics environment to compare expression profiles between (1) cell lines, (2) AAV titer phenotypes, and (3) after transfection timepoints. Unless otherwise noted, we applied a false discovery rate (FDR) threshold of 0.01 and a log_2_ (fold change) threshold of 0.5 to identify top differentially expressed genes across analyses. First, to compare expression profiles across cell lines (Figure 2 and Figure 2 – figure supplement 1), we generated contrasts using pairwise combinations of all 5 cell types (293F, AC1P, AC112, AC203, and AC230) and generated a matrix to compare the number of differentially expressed genes between each pair (Figure 2 – figure supplement 1D). We then performed differential expression tests between the base and high AAV producing groups. Finally, to identify genes which were significantly up- or downregulated during AAV production, we performed pairwise differential expression tests between the 0, 24, and 72 hr time points.

### Gene ontology and drug interaction analysis

To identify sets of coregulated genes associated with high AAV-producing cells, we performed Reactome pathway analysis using the top differentially expressed genes between base and high producing cell lines (Milacic et al., 2024). We identified overrepresented pathways by using a p-value threshold of 0.05. We then manually curated the results for heatmap visualization by combining overlapping pathways into broader categories and removing duplicate genes across categories (Figure 2D). We carried out a similar approach to classify differentially modulated genes during the AAV production time course. Additionally, Gene Ontology (GO) analysis was performed using the BigOmics platform to identify significantly enriched cellular components and molecular functions during AAV production (Figure 4).

We performed L1000 drug connectivity analysis within the BigOmics platform using differentially expressed genes from the 0 vs. 24 hr and the 0 vs. 72 hr timepoint contrasts (Subramanian et al., 2017). Normalized enrichment scores (NES) with p-values <0.01 were used to identify enriched and depleted targets, mechanisms of action, and drug candidates at 24 and 72 hr after transfection.

### Compound screening

Compounds were solubilized in DMSO at a 10 mM stock concentration, and later diluted to the appropriate concentration (0.1-1.0 μM in 0.1% DMSO). 0.1% DMSO was used as the control condition. Microplates or shake flasks were then incubated at 37°C and 5% CO_2_ for 72 hr with no media change prior to sample harvest.

### Statistical analysis

The number of replicates is indicated in the figure legends. Unless otherwise specified, Student’s *t*-test was used for statistical analysis, with significant differences defines as \**P* <0.05, ** *P* <0.01, *** *P*<0.001, **** *P*<0.0001. Error bars indicate standard deviation (SD).

## Data availability statement

Our RNA-Seq data has been deposited on the Gene Expression Omnibus (GEO) database: GEO Submission (GSE269485).

## Acknowledgments

We thank C.R. Herron for editing the manuscript. We thank H. Griffiths and R. Feitzinger, L. Leveque-Eichhorn and I. Ukani for assay optimization and plasmid cloning.

## Author contributions

F.G., C.A.R., and M.A.M. were responsible for the conception and design of the experiments. F.G., and J.T. conducted the experiments. J.T., C.A.R., and M.A.M. performed seq analysis and interpreted the results. C.A.R., M.H., and M.A.M. supervised the studies. J.T., and M.A.M. wrote the manuscript with support from all authors.

## Key resources table

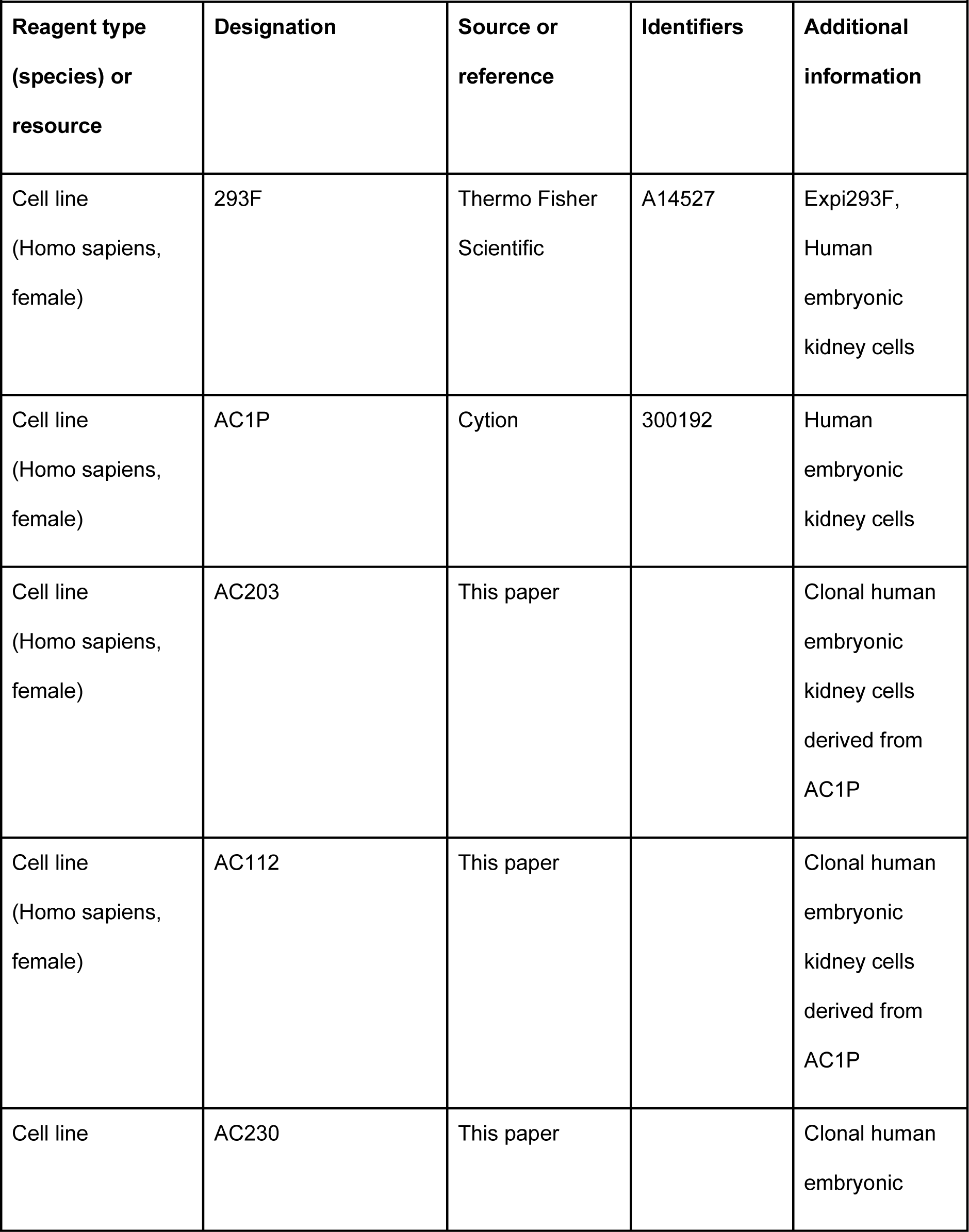
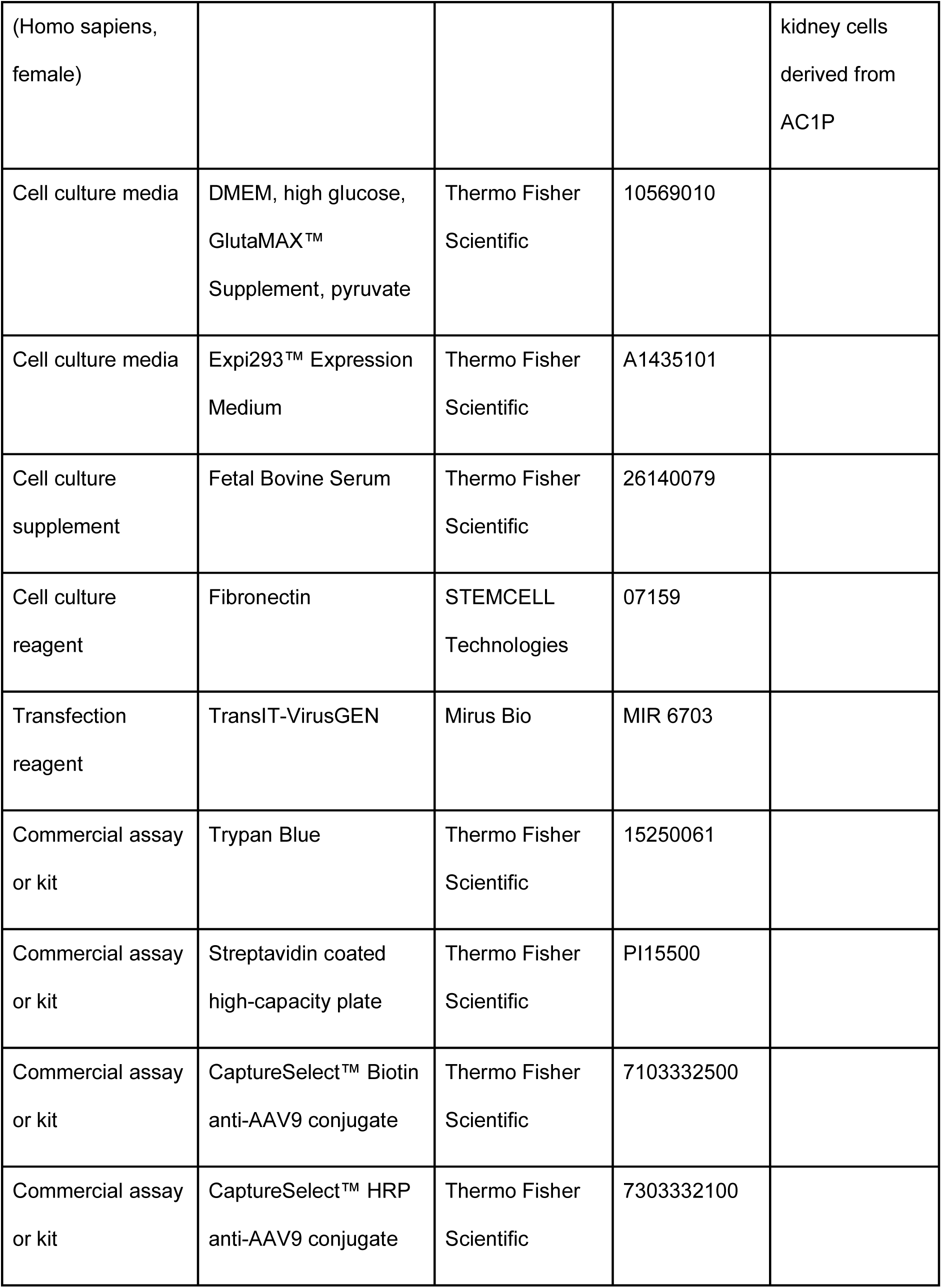
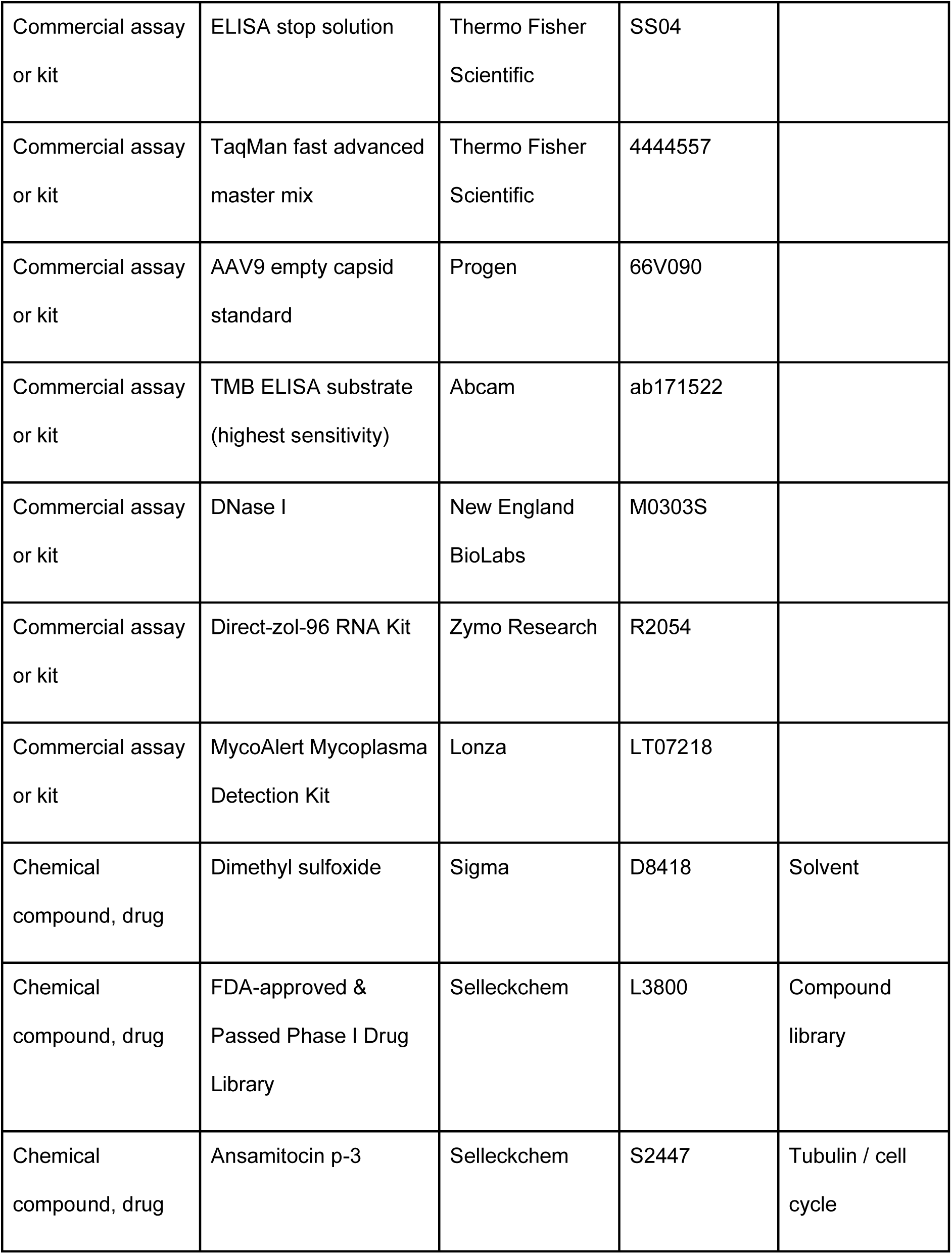
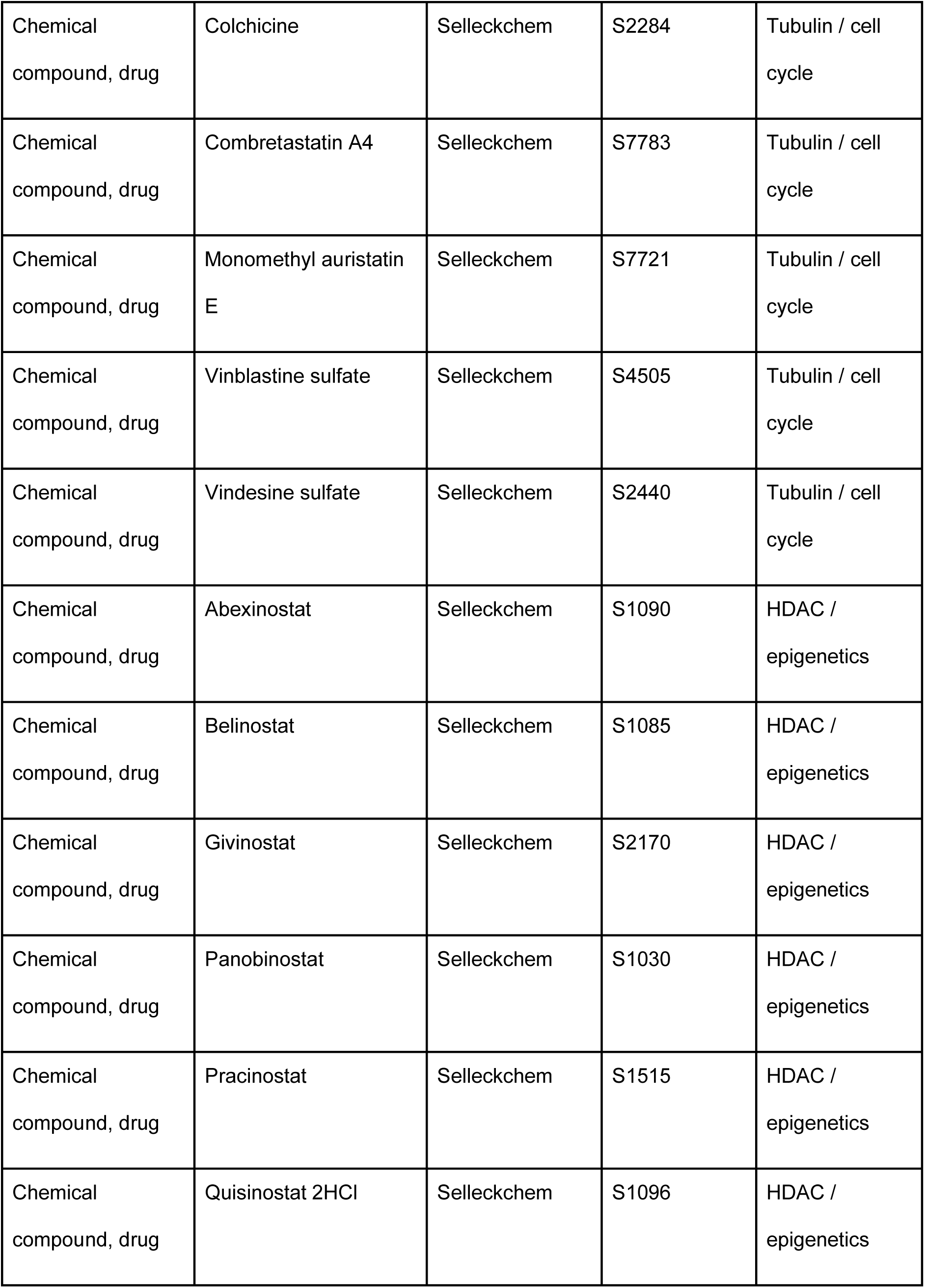
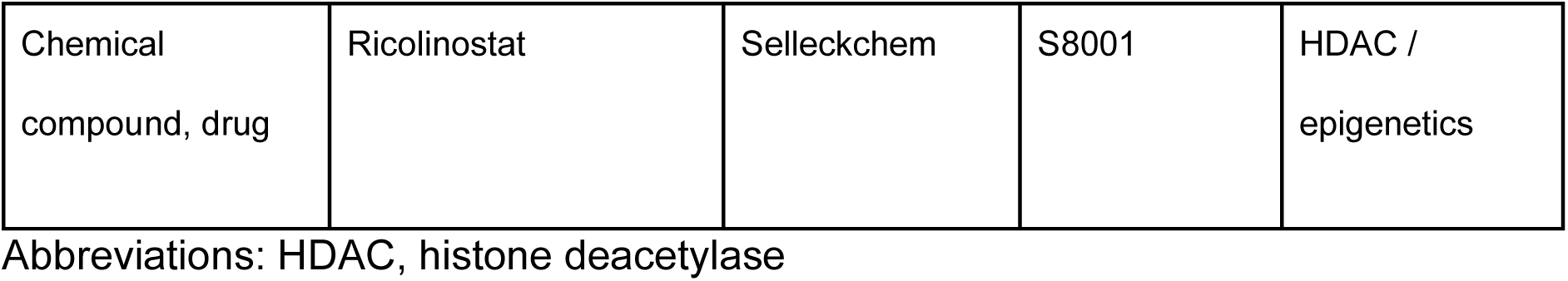

## Supplementary material

**Figure 1 - figure supplement 1.**
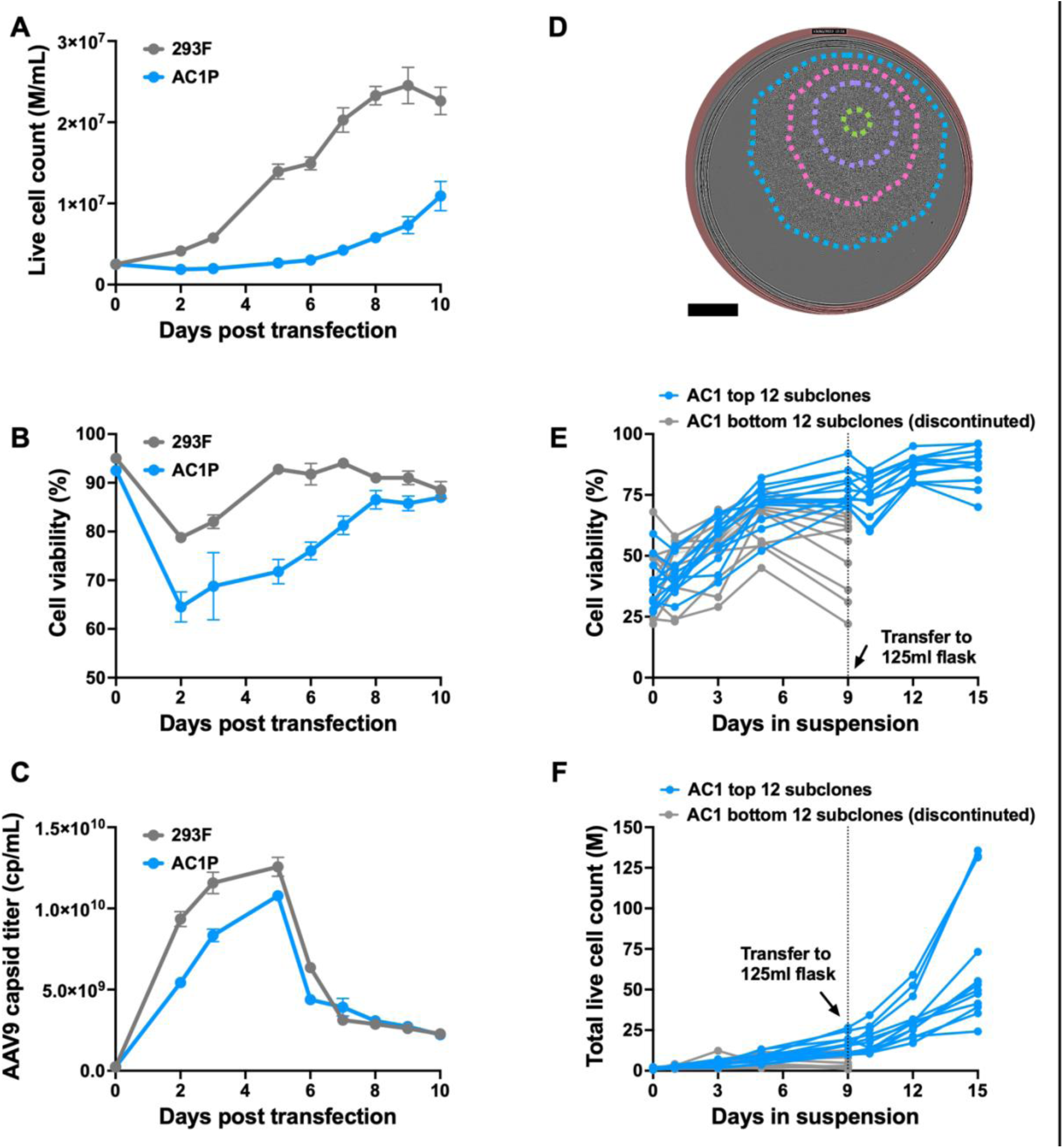
AAV production in 293F and AC1P polyclonal line, and suspension adaptation of AC1P subclones. (**A**) Live cell count, (**B**) cell viability and (**C**) AAV9 capsid titer was measured for 10 days after triple transfection in 293F and AC1P. n = 3 replicates per sample. Data are shown as mean ± SD. (**C**) Representative single-cell derived clonal colony in a 96-well plate. Dashed lines show colony boundary at 3 (green), 5 (purple), 7 (pink), and 10 (blue) days after single-cell seeding. Scale bar, 1000 µm. (**D**) Cell viability and (**F**) live cell count of top 24 clones was measured during the suspension adaptation phase. After 9 days of suspension adaptation, the top 12 clones (based on viability and cell count) were selected and progressed to 125mL shake flasks.

**Figure 2 - figure supplement 1.**
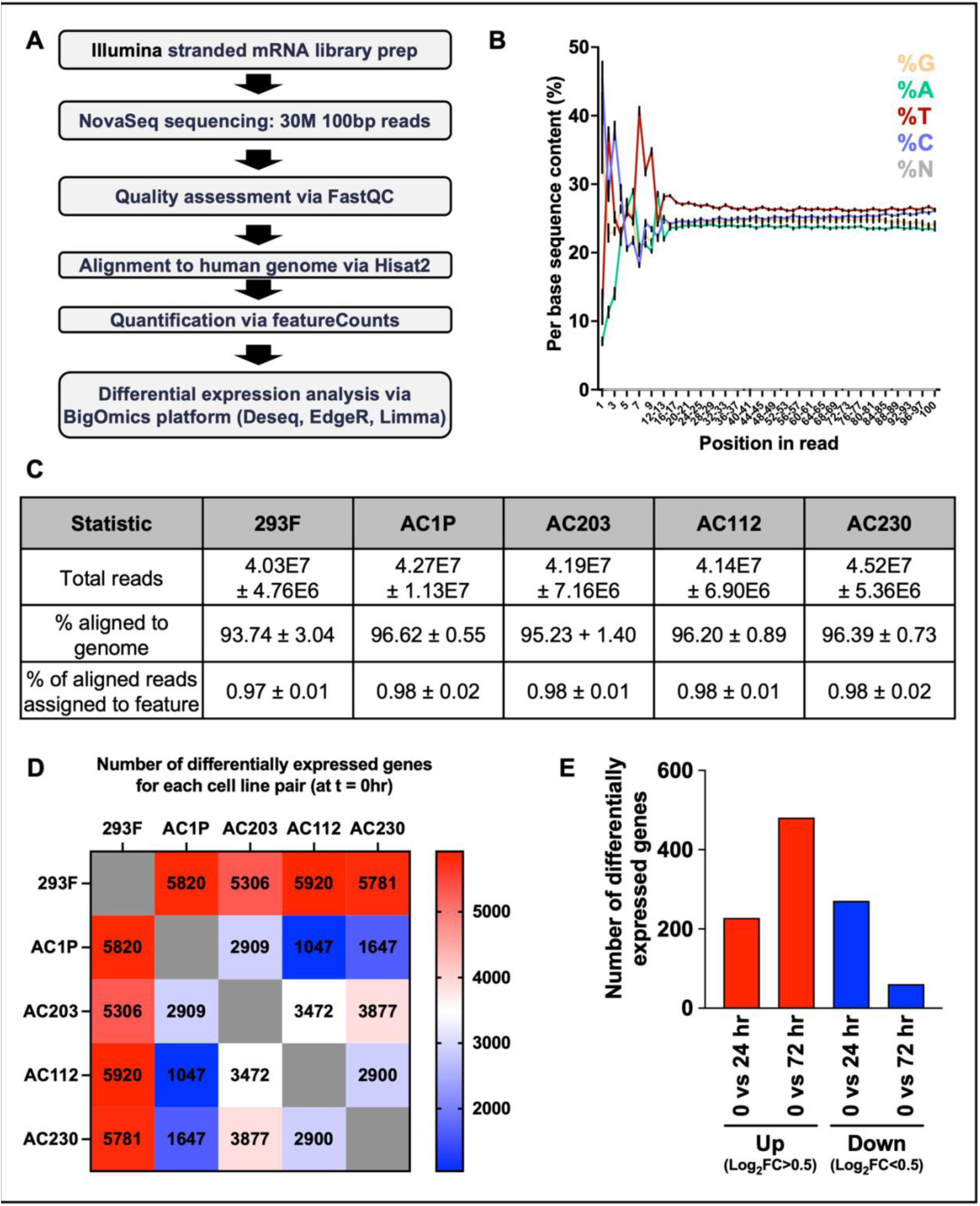
Overview of bioinformatic workflow, RNA-seq quality control and differentially expressed genes between cell lines and across timepoints. (**A**) Bioinformatic workflow for analysis of RNA-seq samples. (**B**) Per-base sequence content across all samples, measured by FastQC. (**C**) Tabulated total reads and sequence alignment results for all samples. (**D**) Pairwise comparison of number of differentially expressed genes at 0hr. (**E**) Number of differentially expressed genes at 24 and 72 hr after transfection (log_2_ (fold change) >0.5 or log_2_ (fold change) <0.5). FC, fold change compared to 0 hr.

**Figure 3 – figure supplement 1.**
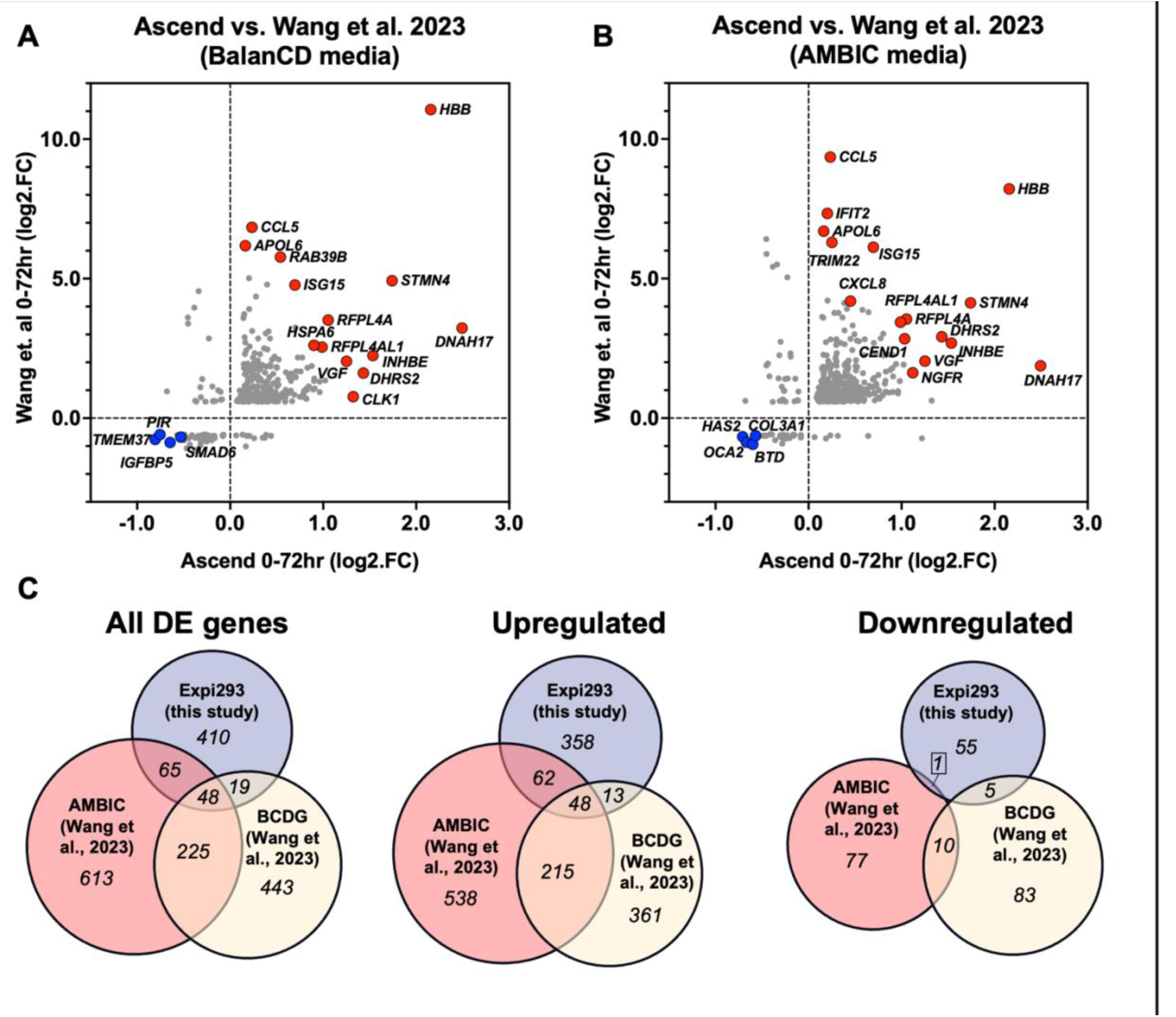
Comparison of transcriptional changes during AAV production to Wang et al. 2023. (**A, B**) Scatter plot comparing transcript fold change among commonly identified genes in current study compared with Wang et al. 2023 in BalanCD (**A**) and AMBIC (**B**) media. A selection of common differentially expressed transcripts during AAV production are denoted in red (upregulated) blue (downregulated) and annotated. For this comparison, we included all differentially expressed transcripts from our study with FDR < 0.05, irrespective of log_2_ (fold change). (**C**) Pie chart comparing the number of shared and unique differentially expressed transcripts in each comparison group. For this comparison, we included only those differentially expressed transcripts with FDR < 0.05 and |log_2_ (fold change)| ≥ 0.5.

**Figure 3 – figure supplement 2.**
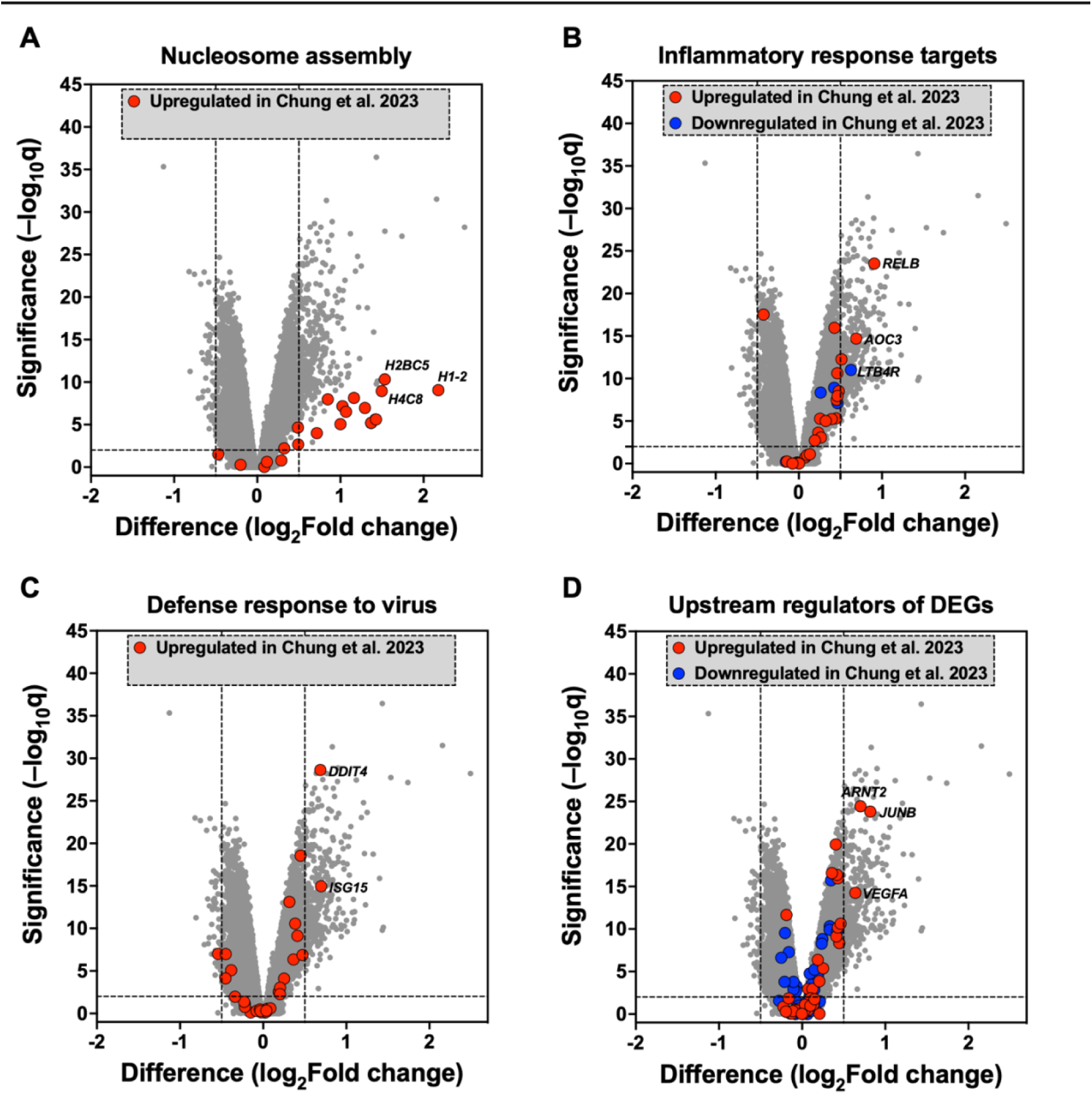
Comparison of transcriptional changes during AAV production to Chung et al. 2023. Differentially expressed genes reported by Chung et al. (2023) was compared to our data and plotted as volcano plots. (**A**) Similar to the Chung et al. 2023 study we see elevation of nucleosome assembly genes at 72 hr. (**B**) Elevation of some inflammatory response and (**C**) virus defense response genes were also detected at 72 hr in our dataset. However, based on gene enrichment analysis, we did not find the majority of the pathways reported by Chung et al. (2023) to be significantly enriched in our dataset. (**D**) Significant activation of a subset of upstream regulators of DEGs reported by Chung et al. (2023) was detected in our data set. Dashed lines on the x axis represent a |log_2_ (fold change)| ≥ 0.5 and the dashed line on the y axis represents a *p*=0.01.

**Figure 4 – figure supplement 1.**
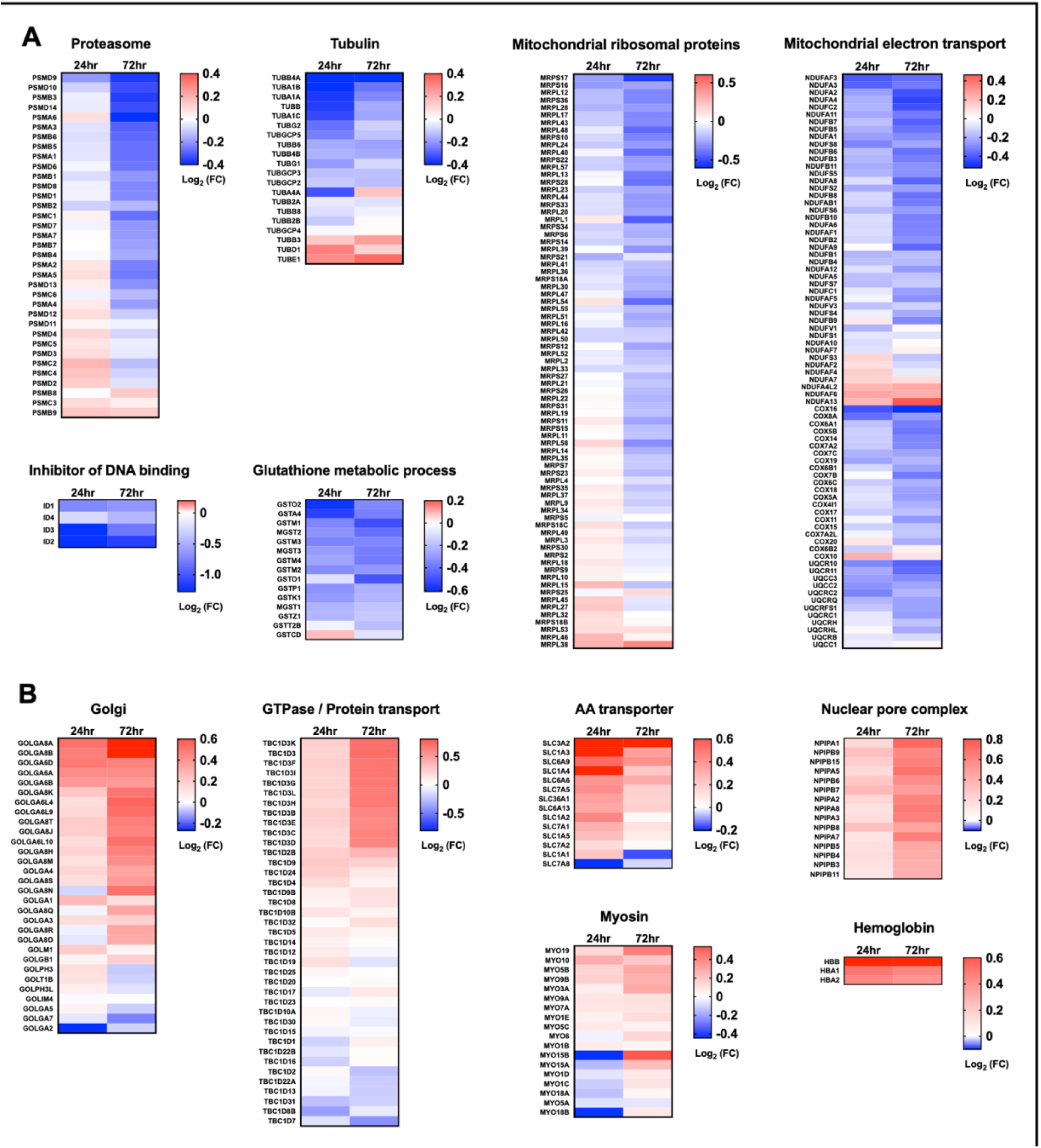
Log_2_ (fold change) heatmaps of differentially expressed genes highlighted among GO terms identified in Figure 4. Heatmaps of genes represented in the GO terms which were significantly (**A**) downregulated and (**B**) upregulated during AAV production. FC, fold change compared to 0 hr.

**Figure 5 – figure supplement 1.**
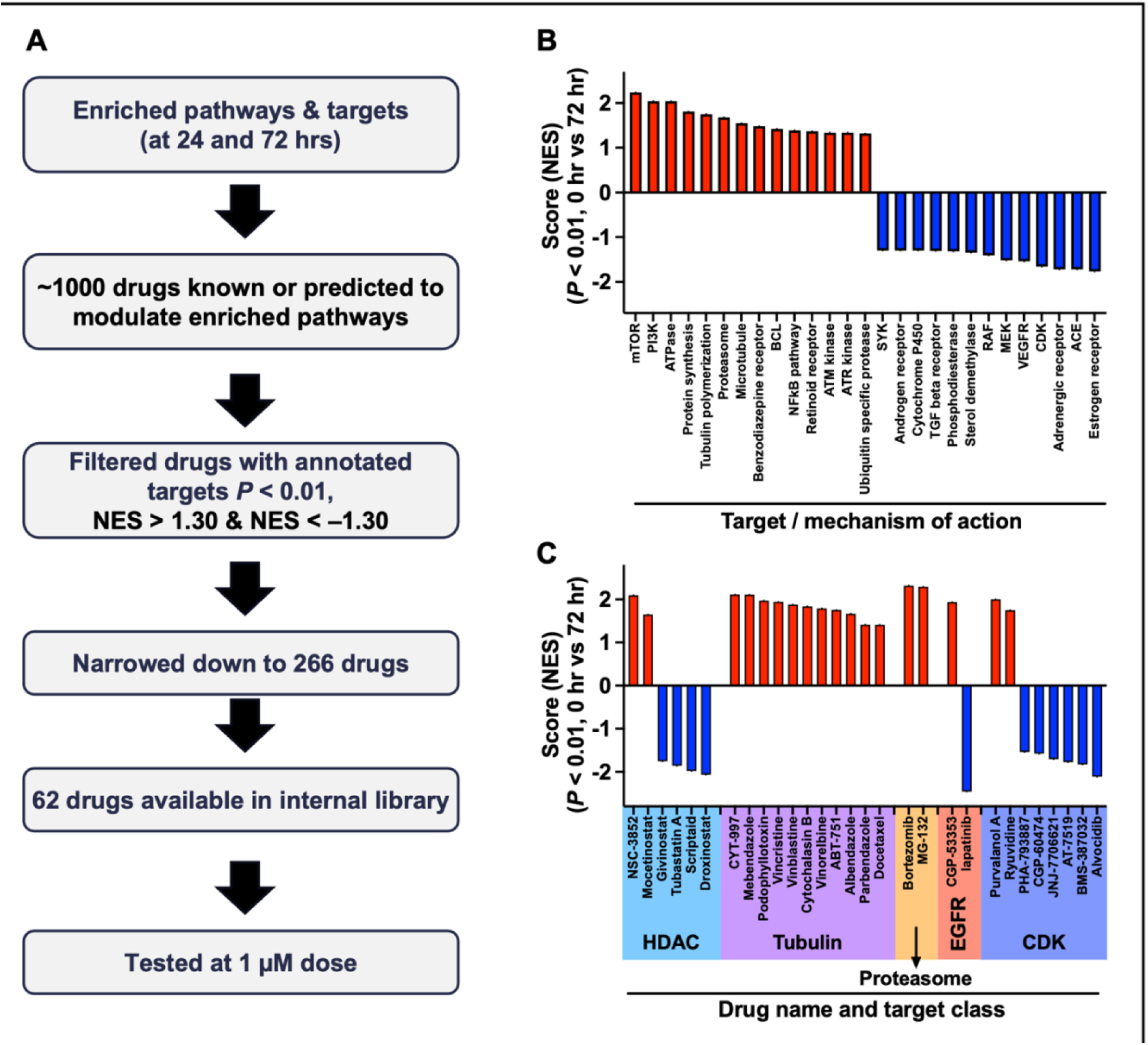
Pathway analysis identifies druggable pathways that may enhance AAV production. (**A**) Workflow and triaging strategy to identify drugs that may enhance AAV production. We identified enriched or depleted pathways at 24 hr (Figure 5) and 72 hr during AAV production. ∼1000 compounds were predicted to modulate the enriched pathways. We next filtered this primary list to only include annotated compounds with known or putative biological targets with a NES greater than 1.3 or less than **–**1.3. Using this criteria, 266 drugs were included in the final list. From the 266 drugs, 62 were present in our screening library and were tested at 1µM dose. (**B**) Normalized enrichment score (NES) with P-values <0.01 was used to identify enriched and depleted targets or mechanisms of action at the late stage of AAV production (0 vs 72 hr). Similar to the 24 hr timepoint, examples of enriched pathways included protein synthesis and tubulin polymerization, while depleted pathways included various class of kinases (mostly involved in cell cycle and cell signaling). (**C**) Normalized enrichment score (NES) with P-values <0.01 among drugs with known targets affecting enriched and depleted pathways at the late stage of AAV production (0 vs 72 hr). Similar to the 24 hr timepoint, enriched drug classes include HDAC, EGFR, Proteasome, and cyclin dependent kinase modulators.

## References

Adachi Y, Yamamoto K, Okada T, Yoshida H, Harada A, Mori K. 2008. ATF6 is a transcription factor specializing in the regulation of quality control proteins in the endoplasmic reticulum. Cell Struct Funct 33:75–89. doi:10.1247/csf.07044

Akhmedov M, Martinelli A, Geiger R, Kwee I. 2020. Omics Playground: a comprehensive self-service platform for visualization, analytics and exploration of Big Omics Data. NAR Genomics and Bioinformatics 2:lqz019. doi:10.1093/nargab/lqz019

Andrews S. 2010. FastQC: A Quality Control Tool for High Throughput Sequence Data [Online]. Available online at: http://www.bioinformatics.babraham.ac.uk/projects/fastqc/.

Bai S, Shi X, Yang X, Cao X. 2000. Smad6 as a transcriptional corepressor. J Biol Chem 275:8267–8270. doi:10.1074/jbc.275.12.8267

Balakrishnan B, Sen D, Hareendran S, Roshini V, David S, Srivastava A, Jayandharan GR. 2013. Activation of the Cellular Unfolded Protein Response by Recombinant Adeno-Associated Virus Vectors. PLoS One 8:e53845. doi:10.1371/journal.pone.0053845

Barnes CR, Lee H, Ojala DS, Lewis KK, Limsirichai P, Schaffer DV. 2021. Genome-wide activation screens to increase adeno-associated virus production. Molecular Therapy - Nucleic Acids 26:94–103. doi:10.1016/j.omtn.2021.06.026

Bennett A, Mietzsch M, Agbandje-McKenna M. 2017. Understanding capsid assembly and genome packaging for adeno-associated viruses. Future Virol 12:283–297. doi:10.2217/fvl-2017-0011

Berthet C, Raj K, Saudan P, Beard P. 2005. How adeno-associated virus Rep78 protein arrests cells completely in S phase. Proceedings of the National Academy of Sciences 102:13634–13639. doi:10.1073/pnas.0504583102

Chung C-H, Murphy CM, Wingate VP, Pavlicek JW, Nakashima R, Wei W, McCarty D, Rabinowitz J, Barton E. 2023. Production of rAAV By Plasmid Transfection Induces Antiviral and Inflammatory Responses in Suspension HEK293 Cells. Molecular Therapy - Methods & Clinical Development 0. doi:10.1016/j.omtm.2023.01.002

Coplan L, Zhang Z, Ragone N, Reeves J, Rodriguez A, Shevade A, Bak H, Tustian AD. 2024. High-yield recombinant adeno-associated viral vector production by multivariate optimization of bioprocess and transfection conditions. Biotechnol Prog e3445. doi:10.1002/btpr.3445

DeLisle AJ, Graves RA, Marzluff WF, Johnson LF. 1983. Regulation of Histone mRNA Production and Stability in Serum-Stimulated Mouse 3T6 Fibroblasts. Molecular and Cellular Biology 3:1920–1929. doi:10.1128/mcb.3.11.1920-1929.1983

Dreiman GHS, Bictash M, Fish PV, Griffin L, Svensson F. 2021. Changing the HTS Paradigm: AI-Driven Iterative Screening for Hit Finding. SLAS Discov 26:257–262. doi:10.1177/2472555220949495

Dumont J, Euwart D, Mei B, Estes S, Kshirsagar R. 2016. Human cell lines for biopharmaceutical manufacturing: history, status, and future perspectives. Crit Rev Biotechnol 36:1110–1122. doi:10.3109/07388551.2015.1084266

Espinoza JA, González PA, Kalergis AM. 2017. Modulation of Antiviral Immunity by Heme Oxygenase-1. The American Journal of Pathology 187:487–493. doi:10.1016/j.ajpath.2016.11.011

FDA. 2024. https://www.fda.gov/. https://www.fda.gov/

Ferreira LT, Figueiredo AC, Orr B, Lopes D, Maiato H. 2018. Dissecting the role of the tubulin code in mitosis. Methods Cell Biol 144:33–74. doi:10.1016/bs.mcb.2018.03.040

Ferreira TB, Perdigão R, Silva AC, Zhang C, Aunins JG, Carrondo MJT, Alves PM. 2009. 293 cell cycle synchronisation adenovirus vector production. Biotechnology Progress 25:235–243. doi:10.1002/btpr.64

Foo J, Bellot G, Pervaiz S, Alonso S. 2022. Mitochondria-mediated oxidative stress during viral infection. Trends in Microbiology 30:679–692. doi:10.1016/j.tim.2021.12.011

Franzoso FD, Seyffert M, Vogel R, Yakimovich A, de Andrade Pereira B, Meier AF, Sutter SO, Tobler K, Vogt B, Greber UF, Büning H, Ackermann M, Fraefel C. 2017. Cell Cycle-Dependent Expression of Adeno-Associated Virus 2 (AAV2) Rep in Coinfections with Herpes Simplex Virus 1 (HSV-1) Gives Rise to a Mosaic of Cells Replicating either AAV2 or HSV-1. J Virol 91:e00357–17. doi:10.1128/JVI.00357-17

Gallinari P, Marco SD, Jones P, Pallaoro M, Steinkühler C. 2007. HDACs, histone deacetylation and gene transcription: from molecular biology to cancer therapeutics. Cell Research 17:195–211. doi:10.1038/sj.cr.7310149

Galluzzi L, Vitale I, Aaronson SA, Abrams JM, Adam D, Agostinis P, Kroemer G, et al. 2018. Molecular mechanisms of cell death: recommendations of the Nomenclature Committee on Cell Death 2018. Cell Death Differ 25:486–541. doi:10.1038/s41418-017-0012-4

Gano JJ, Simon JA. 2010. A Proteomic Investigation of Ligand-dependent HSP90 Complexes Reveals CHORDC1 as a Novel ADP-dependent HSP90-interacting Protein. Mol Cell Proteomics 9:255–270. doi:10.1074/mcp.M900261-MCP200

Grafton F, Feitzinger R, Fisher K, Leveque-Eichhorn L, Reid CA, Mandegar MA. 2023. A High-Throughput Small Molecule Screen Identifies Targets That Increase AAV9 Production in Suspension HEK293 Cells (ASGCT abstract 921). Molecular Therapy Vol 31 No 4S1.

Hart LS, Ornelles D, Koumenis C. 2007. The Adenoviral E4orf6 Protein Induces Atypical Apoptosis in Response to DNA Damage*. Journal of Biological Chemistry 282:6061–6067. doi:10.1074/jbc.M610405200

Higashino F, Yoshida K, Fujinaga Y, Kamio K, Fujinaga K. 1993. Isolation of a cDNA encoding the adenovirus E1A enhancer binding protein: a new human member of the ets oncogene family. Nucleic Acids Res 21:547–553. doi:10.1093/nar/21.3.547

Imamura T, Takase M, Nishihara A, Oeda E, Hanai J, Kawabata M, Miyazono K. 1997. Smad6 inhibits signalling by the TGF-β superfamily. Nature 389:622–626. doi:10.1038/39355

Junod SL, Saredy J, Yang W. 2021. Nuclear Import of Adeno-Associated Viruses Imaged by High-Speed Single-Molecule Microscopy. Viruses 13:167. doi:10.3390/v13020167

Kahlig C-I, Moser S, Micutkova L, Grillari J, Kraus B, Hernandez Bort JA. 2024. Enhancement of rAAV titers via inhibition of the interferon signaling cascade in transfected HEK293 suspension cultures. Biotechnol J 19:e2300672. doi:10.1002/biot.202300672

Khan N, Cheemadan S, Saxena H, Bammidi S, Jayandharan GR. 2020. MicroRNA-based recombinant AAV vector assembly improves efficiency of suicide gene transfer in a murine model of lymphoma. Cancer Medicine 9:3188–3201. doi:10.1002/cam4.2935

Kim D, Paggi JM, Park C, Bennett C, Salzberg SL. 2019. Graph-based genome alignment and genotyping with HISAT2 and HISAT-genotype. Nat Biotechnol 37:907–915. doi:10.1038/s41587-019-0201-4

Kuzmin DA, Shutova MV, Johnston NR, Smith OP, Fedorin VV, Kukushkin YS, van der Loo JCM, Johnstone EC. 2021. The clinical landscape for AAV gene therapies. Nat Rev Drug Discov 20:173–174. doi:10.1038/d41573-021-00017-7

Lasorella A, Benezra R, Iavarone A. 2014. The ID proteins: master regulators of cancer stem cells and tumour aggressiveness. Nat Rev Cancer 14:77–91. doi:10.1038/nrc3638

Liao Y, Smyth GK, Shi W. 2014. featureCounts: an efficient general purpose program for assigning sequence reads to genomic features. Bioinformatics 30:923–930. doi:10.1093/bioinformatics/btt656

Lipsick JS, Baluda MA. 1986. The myb oncogene. Gene Amplif Anal 4:73–98.

Liu W, Baker SS, Baker RD, Nowak NJ, Zhu L. 2011. Upregulation of Hemoglobin Expression by Oxidative Stress in Hepatocytes and Its Implication in Nonalcoholic Steatohepatitis. PLoS One 6:e24363. doi:10.1371/journal.pone.0024363

Milacic M, Beavers D, Conley P, Gong C, Gillespie M, Griss J, Haw R, Jassal B, Matthews L, May B, Petryszak R, Ragueneau E, Rothfels K, Sevilla C, Shamovsky V, Stephan R, Tiwari K, Varusai T, Weiser J, Wright A, Wu G, Stein L, Hermjakob H, D’Eustachio P. 2024. The Reactome Pathway Knowledgebase 2024. Nucleic Acids Research 52:D672–D678. doi:10.1093/nar/gkad1025

Nakabeppu Y, Nathans D. 1991. A naturally occurring truncated form of FosB that inhibits Fos/Jun transcriptional activity. Cell 64:751–759. doi:10.1016/0092-8674(91)90504-R

Nguyen TNT, Sha S, Hong MS, Maloney AJ, Barone PW, Neufeld C, Wolfrum J, Springs SL, Sinskey AJ, Braatz RD. 2021. Mechanistic model for production of recombinant adeno-associated virus via triple transfection of HEK293 cells. Molecular Therapy - Methods & Clinical Development 21:642–655. doi:10.1016/j.omtm.2021.04.006

Nicolson SC, Samulski RJ. 2014. Recombinant Adeno-Associated Virus Utilizes Host Cell Nuclear Import Machinery To Enter the Nucleus. J Virol 88:4132–4144. doi:10.1128/JVI.02660-13

Onishi T, Nonaka M, Maruno T, Yamaguchi Y, Fukuhara M, Torisu T, Maeda M, Abbatiello S, Haris A, Richardson K, Giles K, Preece S, Yamano-Adachi N, Omasa T, Uchiyama S. 2023. Enhancement of recombinant adeno-associated virus activity by improved stoichiometry and homogeneity of capsid protein assembly. Molecular Therapy Methods & Clinical Development 31. doi:10.1016/j.omtm.2023.101142

Paszkiet BJ, Zhang J, Matukonis M, Kaleko M, Luo T. 2022. Histone Deacetylation Inhibitors Enhance Lentiviral Vector Production and Infectivity (Abstract 944). Molecular Therapy **Vol.** 5**, No.** 5:S308. doi:10.1016/S1525-0016(16)43774-3

Peng Y, Kang Q, Luo Q, Jiang W, Si W, Liu BA, Luu HH, Park JK, Li X, Luo J, Montag AG, Haydon RC, He T-C. 2004. Inhibitor of DNA Binding/Differentiation Helix-Loop-Helix Proteins Mediate Bone Morphogenetic Protein-induced Osteoblast Differentiation of Mesenchymal Stem Cells. Journal of Biological Chemistry 279:32941–32949. doi:10.1074/jbc.M403344200

Polo SE, Almouzni G. 2005. Histone metabolic pathways and chromatin assembly factors as proliferation markers. Cancer Letters 220:1–9. doi:10.1016/j.canlet.2004.08.024

Pupo A, Fernández A, Low SH, François A, Suárez-Amarán L, Samulski RJ. 2022. AAV vectors: The Rubik’s cube of human gene therapy. Molecular Therapy 30:3515–3541. doi:10.1016/j.ymthe.2022.09.015

Ryseck RP, Bull P, Takamiya M, Bours V, Siebenlist U, Dobrzanski P, Bravo R. 1992. RelB, a new Rel family transcription activator that can interact with p50-NF-kappa B. Mol Cell Biol 12:674–684. doi:10.1128/mcb.12.2.674-684.1992

Samulski RJ, Muzyczka N. 2014. AAV-Mediated Gene Therapy for Research and Therapeutic Purposes. Annu Rev Virol 1:427–451. doi:10.1146/annurev-virology-031413-085355

Scarrott JM, Johari YB, Pohle TH, Liu P, Mayer A, James DC. 2023. Increased recombinant adeno-associated virus production by HEK293 cells using small molecule chemical additives. Biotechnology Journal 18:2200450. doi:10.1002/biot.202200450

Schweitzer-Stenner R. 2022. Heme–Protein Interactions and Functional Relevant Heme Deformations: The Cytochrome c Case. Molecules 27. doi:10.3390/molecules27248751

Shahid-Fuente IW, Toseland CP. 2023. Myosin in chromosome organisation and gene expression. Biochemical Society Transactions 51:1023–1034. doi:10.1042/BST20220939

Sieweke MH, Tekotte H, Frampton J, Graf T. 1997. MafB represses erythroid genes and differentiation through direct interaction with c-Ets-1. Leukemia 11 **Suppl 3**:486–488.

Srivastava A, Mallela KMG, Deorkar N, Brophy G. 2021. Manufacturing Challenges and Rational Formulation Development for AAV Viral Vectors. J Pharm Sci 110:2609–2624. doi:10.1016/j.xphs.2021.03.024

Subramanian A, Narayan R, Corsello SM, Peck DD, Natoli TE, Lu X, Gould J, Davis JF, Tubelli AA, Asiedu JK, Lahr DL, Hirschman JE, Liu Z, Donahue M, Julian B, Khan M, Wadden D, Smith I, Lam D, Liberzon A, Toder C, Bagul M, Orzechowski M, Enache OM, Piccioni F, Johnson SA, Lyons NJ, Berger AH, Shamji A, Brooks AN, Vrcic A, Flynn C, Rosains J, Takeda D, Hu R, Davison D, Lamb J, Ardlie K, Hogstrom L, Greenside P, Gray NS, Clemons PA, Silver S, Wu Xiaoyun, Zhao W-N, Read-Button W, Wu Xiaohua, Haggarty SJ, Ronco LV, Boehm JS, Schreiber SL, Doench JG, Bittker JA, Root DE, Wong B, Golub TR. 2017. A Next Generation Connectivity Map: L1000 platform and the first 1,000,000 profiles. Cell 171:1437–1452.e17. doi:10.1016/j.cell.2017.10.049

Tan E, Chin CSH, Lim ZFS, Ng SK. 2021. HEK293 Cell Line as a Platform to Produce Recombinant Proteins and Viral Vectors. Front Bioeng Biotechnol 9. doi:10.3389/fbioe.2021.796991

Turner-Gillies E, Shapiro B, Tian F. 2024. Generation of Cell Lines Capable of Producing High-titer Viral Stocks for Use in Vaccine Manufacture and Gene Therapy. ATCC Application note.

Wada R. 2022. Enhancement of rAAV Productivity Utilizing HDAC Inhibitors on Helper-Free HEK293 Suspension Cell Culture Process (ASGCT abstract 766). Molecular Therapy Vol 30 No 4S1.

Wang J-H, Gessler DJ, Zhan W, Gallagher TL, Gao G. 2024. Adeno-associated virus as a delivery vector for gene therapy of human diseases. Sig Transduct Target Ther 9:1–33. doi:10.1038/s41392-024-01780-w

Wang Y, Fu Q, Lee YS, Sha S, Yoon S. 2023. Transcriptomic features reveal molecular signatures associated with recombinant adeno-associated virus production in HEK293 cells. Biotechnology Progress 39:e3346. doi:10.1002/btpr.3346

Wilkinson MG, Millar JB. 2000. Control of the eukaryotic cell cycle by MAP kinase signaling pathways. FASEB J 14:2147–2157. doi:10.1096/fj.00-0102rev

Wright JF. 2023. AAV vector production: Troublesome host innate responses in another setting. Molecular Therapy Methods & Clinical Development 28:412–413. doi:10.1016/j.omtm.2023.02.008

Yang J, Grafton F, Ranjbarvaziri S, Budan A, Farshidfar F, Cho M, Xu E, Ho J, Maddah M, Loewke KE, Medina J, Sperandio D, Patel S, Hoey T, Mandegar MA. 2022. Phenotypic screening with deep learning identifies HDAC6 inhibitors as cardioprotective in a BAG3 mouse model of dilated cardiomyopathy. Science Translational Medicine 14:eabl5654. doi:10.1126/scitranslmed.abl5654

Yang J, Li X, Li Y, Southwood M, Ye L, Long L, Al-Lamki RS, Morrell NW. 2013. Id proteins are critical downstream effectors of BMP signaling in human pulmonary arterial smooth muscle cells. American Journal of Physiology-Lung Cellular and Molecular Physiology 305:L312–L321. doi:10.1152/ajplung.00054.2013

Zhang R, Rapin N, Ying Z, Shklanka E, Bodnarchuk TW, Verge VMK, Misra V. 2013. Zhangfei/CREB-ZF - a potential regulator of the unfolded protein response. PLoS One 8:e77256. doi:10.1371/journal.pone.0077256

Zhang W, Liu HT. 2002. MAPK signal pathways in the regulation of cell proliferation in mammalian cells. Cell Res 12:9–18. doi:10.1038/sj.cr.7290105

Zhang X, Yu W. 2022. Heat shock proteins and viral infection. Front Immunol 13:947789. doi:10.3389/fimmu.2022.947789

